# Metabolic reprogramming tips vaccinia virus infection outcomes by stabilizing interferon-γ induced IRF1

**DOI:** 10.1101/2024.10.10.617691

**Authors:** Tyron Chang, Jessica Alvarez, Sruthi Chappidi, Stacey Crockett, Mahsa Sorouri, Robert C. Orchard, Dustin C. Hancks

## Abstract

Interferon (IFN) induced activities are critical, early determinants of immune responses and infection outcomes. A key facet of IFN responses is the upregulation of hundreds of mRNAs termed interferon-stimulated genes (ISGs) that activate intrinsic and cell-mediated defenses. While primary interferon signaling is well-delineated, other layers of regulation are less explored but implied by aberrant ISG expression signatures in many diseases in the absence of infection. Consistently, our examination of tonic ISG levels across uninfected human tissues and individuals revealed three ISG subclasses. As tissue identity and many comorbidities with increased virus susceptibility are characterized by differences in metabolism, we characterized ISG responses in cells grown in media known to favor either aerobic glycolysis (glucose) or oxidative phosphorylation (galactose supplementation). While these conditions over time had a varying impact on the expression of ISG RNAs, the differences were typically greater between treatments than between glucose/galactose. Interestingly, extended interferon-priming led to divergent expression of two ISG proteins: upregulation of IRF1 in IFN-γ/glucose and increased IFITM3 in galactose by IFN-α and IFN-γ. In agreement with a hardwired response, glucose/galactose regulation of interferon-γ induced IRF1 is conserved in unrelated mouse and cat cell types. In galactose conditions, proteasome inhibition restored interferon-γ induced IRF1 levels to that of glucose/interferon-γ. Glucose/interferon-γ decreased replication of the model poxvirus vaccinia at low MOI and high MOIs. Vaccinia replication was restored by *IRF1* KO. In contrast, but consistent with differential regulation of IRF1 protein by glucose/galactose, WT and *IRF1* KO cells in galactose media supported similar levels of vaccinia replication regardless of IFN-γ priming. Also associated with glucose/galactose is a seemingly second block at a very late stage in viral replication which results in reductions in herpes- and poxvirus titers but not viral protein expression. Collectively, these data illustrate a novel layer of regulation for the key ISG protein, IRF1, mediated by glucose/galactose and imply unappreciated subprograms embedded in the interferon response. In principle, such cellular circuitry could rapidly adapt immune responses by sensing changing metabolite levels consumed during viral replication and cell proliferation.

## INTRODUCTION

The adaptability of immune responses is essential to respond to selective pressure imposed by pathogen infections over evolutionary time [1]. Established evolutionary mechanisms implicated in counteracting pathogen infection include gene expansions, gene loss [2–4], and rapid evolution of protein-coding genes [5]. Adaptive signatures are notably evident in factors important in establishing early host defenses [6]. Interindividual variation in early responses to infection are well-documented. The most dramatic examples are associated with genetic deficiencies for specific antiviral factors [7]. In addition, there is a growing appreciation for non-genetic risk factors in shaping infection outcomes [8,9] but the molecular basis is incompletely understood. While the contribution of host genetics to the spectrum of infection outcomes continues to come into focus, how non-genetic factors perturb immune responses like signaling remains less clear.

Innate immune signaling in vertebrates is composed of two sequential modalities [10,11]. The first involves recognition of pathogen associated molecular patterns (PAMP) such as viral double-stranded DNA (dsDNA) or dsRNA in initially infected cells [12–14]. PAMP detection is mediated by germline-encoded pattern recognition receptors (PRRs). Well-characterized PRRs are Toll-like receptors (TLR) [15] and cyclic GMP-AMP synthase (cGAS) [12,16]. Generally, PRR signaling induces gene expression of essential cytokines like interferons (IFN), which comprise the second modality. IFNs act in an autocrine and paracrine manner to upregulate hundreds of gene products termed interferon-stimulated genes (ISGs); many of which have direct antiviral activity [17].

Different types of IFNs have been identified that have both distinct and overlapping functions in immune and non-immune cells [18,19]. The type I IFN response, which is considered the primary innate immune response during viral infection in mammals, is induced by IFN-α subtypes and IFN-β. Type I IFNs act on both immune and non-immune cells and are commonly studied in macrophages and epithelial cells [20]. The type II IFN response is triggered by IFN-γ. IFN-γ plays a crucial role in the activation of certain immune cells like macrophages and serves as a catalyst of host defenses in non-immune cells [21]. The type III IFN response is regulated primarily by IFN-λ1, IFN-λ2, and IFN-λ3. IFN-λ signaling has emerged as an essential antiviral response [22] that is mostly limited to cells present at mucosal barrier surfaces in the respiratory and gastrointestinal tract.

Upon IFN receptor ligation, the induction of IFN signaling is mediated by Janus kinase (JAK) and signal transducer and activator of transcription (STAT) proteins [23,24]. However, the molecular components vary depending on the type of IFN binding to the receptor. Aberrant interferon signaling is a hallmark of monogenic diseases - the interferonopathies [25,26] - as well as complex diseases like lupus and diabetes [27,28]. The loss-of-function of specific ISGs linked to poor infection outcomes [29] implies variable expression of specific ISGs may contribute to the heterogeneity of antiviral responses. Dysregulated ISG signatures in the absence of infection foreshadow unappreciated regulatory mechanisms of interferon and ISG responses. For instance, metabolic disorders like mitochondrial diseases are considered comorbidities that display ISG as well as inflammatory gene signatures [30]. Many of these conditions also display increased susceptibility to virus associated disease [8]. Altogether these data suggest ill-defined roles for metabolites and mitochondrial activities in regulating early immune responses linked to infection outcomes.

While IFNs are known to be key drivers of antiviral responses, it is less defined how additional cues shape primary interferon signaling and responses. Recently, lactate has been shown to bind mitochondrial antiviral signaling protein (MAVS) and inhibit its ability to activate type I interferon expression [31]. It is unknown whether other metabolites modulate the magnitude or composition of the ISG response. Changes to the ISG repertoire can tip infection outcomes as indicated by studies of specific host defense proteins inactivated by viral-encoded antagonists [32,33]. However, it is poorly understood whether altering ISG repertoires within a given cell type favors host outcomes. Nevertheless, characterization of new modulators of ISG responses may inform novel regulatory mechanisms for these pivotal defenses and shed light on non-genetic factors contributing to interindividual variation in immune responses.

To gain insights into the regulation of ISG responses, we first characterized the landscape of ISG expression across human tissues at steady state, basal (tonic) levels. In addition to highlighting widespread variation in tonic ISG RNA levels across and within tissues, this analysis revealed three distinct ISG subclasses seemingly independent of interferon and interferon receptor expression. As tissue identity and many comorbidities with increased virus susceptibility are characterized by differences in metabolism, we explored the role of two metabolites on ISG responses: glucose and galactose. We focused on these two metabolites as they are known to tip the cellular balance between glycolysis (glucose) and oxidative phosphorylation (galactose); a phenomenon dysregulated in many co-morbidities. We find that there is time-dependent interplay between glucose/galactose and interferons impacting expression of ISGs. Interestingly, extended interferon priming leads to divergent expression between glucose/galactose of the key ISG and transcription factor - interferon regulatory factor 1 (IRF1). IRF1 is selectively stabilized in IFN-γ treated cells in glucose media, while galactose media promotes its proteasomal degradation. Differential regulation of IRF1 by glucose/galactose is associated with infection outcomes. Specifically, reduced replication of the prototypical poxvirus, vaccinia, in glucose/IFN-γ compared to glucose/untreated and glucose/IFN-α is relieved by *IRF1* KO. In contrast, vaccinia replicates to similar levels in galactose/IFN-γ media regardless of IRF1 status (WT vs. *IRF1* KO cells). These findings illustrate an unappreciated layer of ISG regulation and allude to a cellular program that could act to rapidly adapt ISG responses by sensing the state of infection via consumption of metabolites.

## RESULTS

### Tonic ISG RNAs stratify into three subclasses across human tissues

Recent studies using IFN-treated immune cell types isolated from mice [34] and human embryonic stem cells (ESC) [35] suggest unappreciated variation in expression and regulation of this key class of immune factors. To shed light on potential regulatory mechanisms of ISGs in humans, we assessed expression of ISGs, interferons, and interferon receptors in tissues by exploiting the observation that ISGs are expressed at tonic levels [36]. In this analysis, we were especially interested in barrier tissues (e.g., lung) as they often represent the initial site of infection. Specifically, we analyzed baseline RNA levels for forty-eight ISGs for fifty-four tissues derived from tens to nearly one-thousand cadavers per tissue [37,38]. These forty-eight ISGs were defined as “core ISGs” as their upregulation by type I interferon is conserved in vertebrates [39]. By looking at conserved ISGs in terms of inducibility instead of all ISGs for a given species, which can be species-specific and often arbitrarily defined by fold-change and statistical cut-off, we reasoned we may identify conserved regulons. To view any potential patterns in light of established relationships, we also included expression for thirty-two interferons and six interferon receptors associated with type I, type II, and type III interferon responses. Analysis of median log_2_ transcript per million (TPM) values revealed stark inter-tissue variation in ISG expression (Fig 1, S1 Table). Noticeably, expression of tonic ISGs across tissues clustered into three general subclasses: 1) ubiquitously and constitutively on (high), 2) ubiquitously and constitutively off (low), and 3) differentially expressed (variable). This stratification of ISGs appears independent of IFN expression as these immune signals are expressed at low levels uniformly across all fifty-four tissues examined. The IFN receptors display more variation in tissue expression with four of the receptors belonging to the variable class (*IFNGR1*, *IL10RB*, *IFNAR1*, *IFNAR2*) whereas *IFNGR1* is in the high class and *IFNLR1* is in the low class. These data suggest that there are three tonic ISG RNA subclasses due to a type of regulation independent of canonical interferon signaling and not a common infection because: 1) each profile reflects the median expression for tens to hundreds of samples, 2) interferon expression is low overall for all tissues examined, and 3) ISGs in the low class, like OAS1 and IFIT2, would be expressed at markedly higher levels in infected tissues.

**Fig 1.**
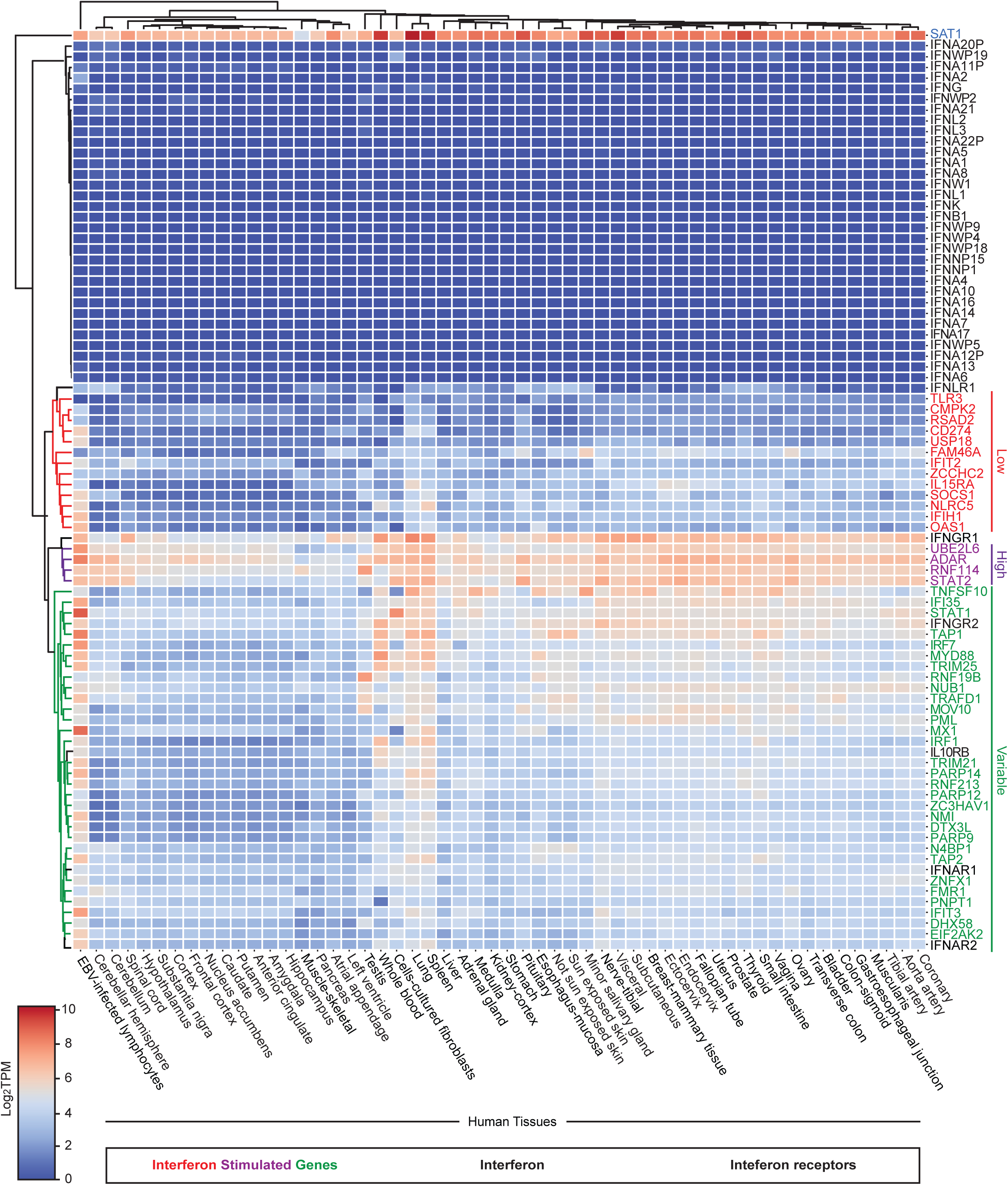
Tonic ISG, interferon, and interferon receptor RNA expression across human tissues. Heatmap displaying gene expression values (median log_2_ TPM) for forty-eight core ISGs, thirty-two human IFNs, and six IFN receptors across fifty-four human tissues. The ISGs cluster into three groups based on tonic expression patterns across tissues: low (red), high (purple), variable (green). Gene names colored in black are either IFNs or IFN receptor genes. *SAT1* (blue) was deemed an outlier following the clustering analysis. TPM: transcripts per million.

To further characterize these tonic ISG expression patterns, we performed Ingenuity Pathway Analysis (IPA) for the three tonic ISG subclasses (S1 Fig, S1 Table). Our analysis found gene ontology terms enriched for each subclass including: low (pathogenesis of influenza, PRR recognition of bacteria and viruses), high (interferon signaling, activation of IRF), and variable (pathogenesis of influenza, retinoic acid mediated apoptosis signaling). The ISGylation signaling pathway was in the top five gene ontology terms for all three subclasses. Next, we exploited the wealth of samples to inspect variation in tonic ISG expression for a subset of tissues that represent key battlegrounds during viral infection. Specifically, we examined representative ISGs from each subclass [high (*ADAR, STAT2*); low (*IFIH1, OAS1*); variable (*IRF1, IRF7, MX1, MYD88, STAT1*)] (Fig 2, S1 Table) for lung, liver, whole blood, and spleen tissues; n = 220 – 755 individual tissue samples. This analysis showed extensive variability for these marker ISGs between individuals and different tissues. For example, *STAT2* is expressed at high levels for all individual samples for lung, liver, and spleen but exhibits a wide range in expression variation in whole blood. Collectively, these findings demonstrate that tonic ISG expression can be divided into three subclasses but is heterogenous across and within tissues as well as individuals. These data suggest poorly characterized layers of ISG regulation including stratification into subclasses.

**Fig 2.**
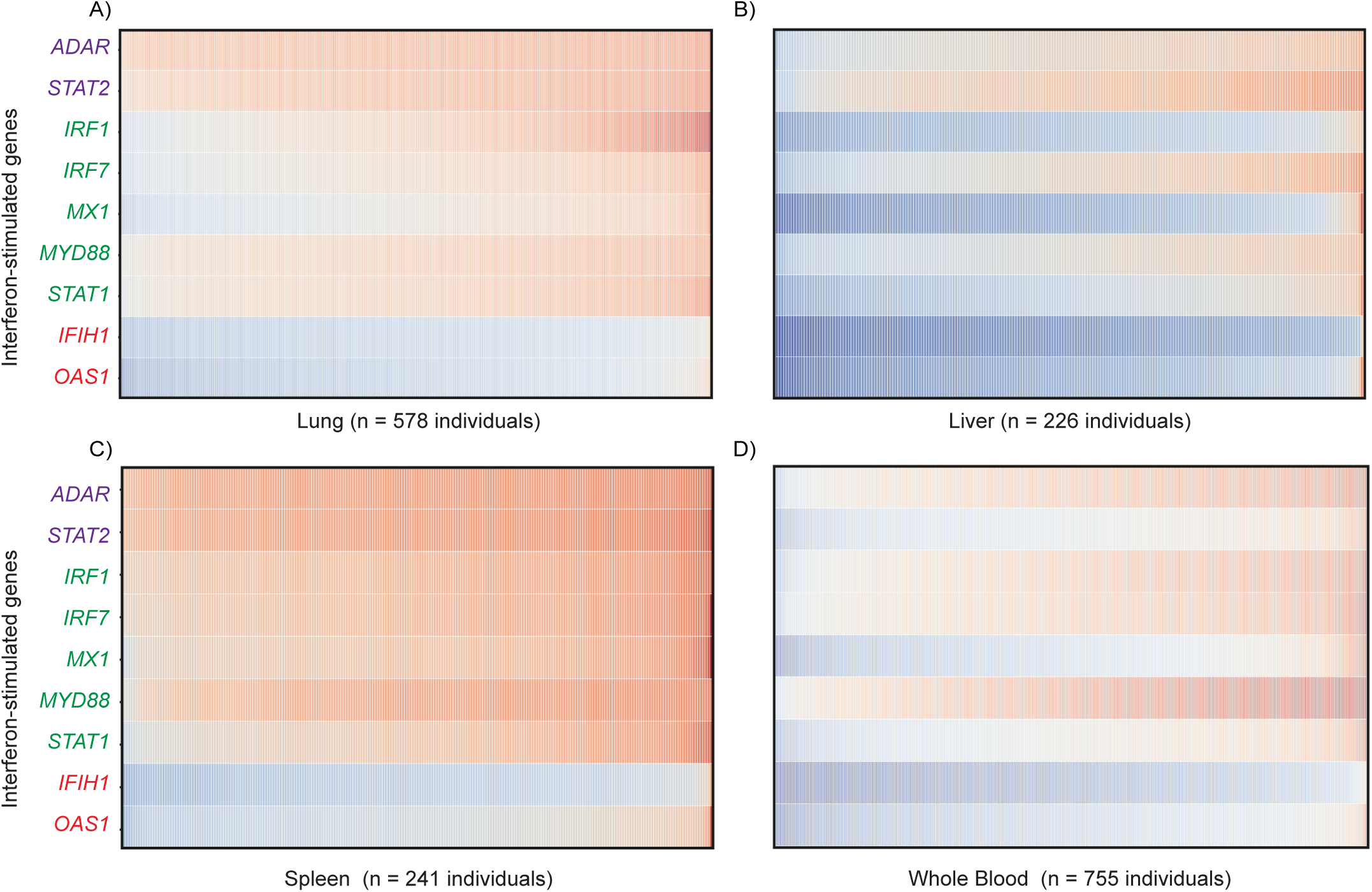
Inter-individual variation in tonic ISG RNA expression in human tissues. RNA expression (log_2_ TPM) of nine core ISGs for (A) 578 individual human lung tissues, (B) 226 human liver tissues, (C) 241 human spleen tissues, and (D) 755 human whole blood tissues. Representative ISGs from each subclass are shown: high (*ADAR, STAT2*), variable (*IRF1*, *IRF7*, *MYD88*, *MX1*, *STAT1*), low (*IFIH1*, *OAS1*). TPM: transcript per million.

### Extended glucose/IFN-γ priming leads to divergent protein expression of specific ISGs

Cell type specific gene expression and activity are influenced by diverse variables [40–42] including metabolism. Changes in cellular metabolism occur as intracellular metabolites are consumed during the activation of host defenses [43] as well as viral replication [44–46] and as a consequence of inflammation [47]. Furthermore, aberrant ISG gene signatures and poor infection outcomes are also associated with certain metabolic diseases [30,48–50] and imply incompletely understood roles for metabolism and/or metabolites in regulating innate immune signaling and host responses.

To examine potential roles for metabolism as a factor influencing the variation in ISG expression highlighted by the tonic ISG analysis, we used a routine strategy to tip cellular metabolism and unmask regulators of mitochondrial activities. The latter of which continue to be linked to immune defense [51–55]. Specifically, we replaced glucose in media, which favors aerobic glycolysis, with galactose, which is known to promote OXPHOS (Fig 3A) [56] and increase sensitivity to mitochondrial toxicants [57]. Similar metabolic reprogramming occurs when macrophages and other immune cells become activated [58,59]. Consistent with increased glycolysis, we observed increased levels of extracellular lactate in glucose media relative to galactose conditions (S2A Fig).

**Fig 3.**
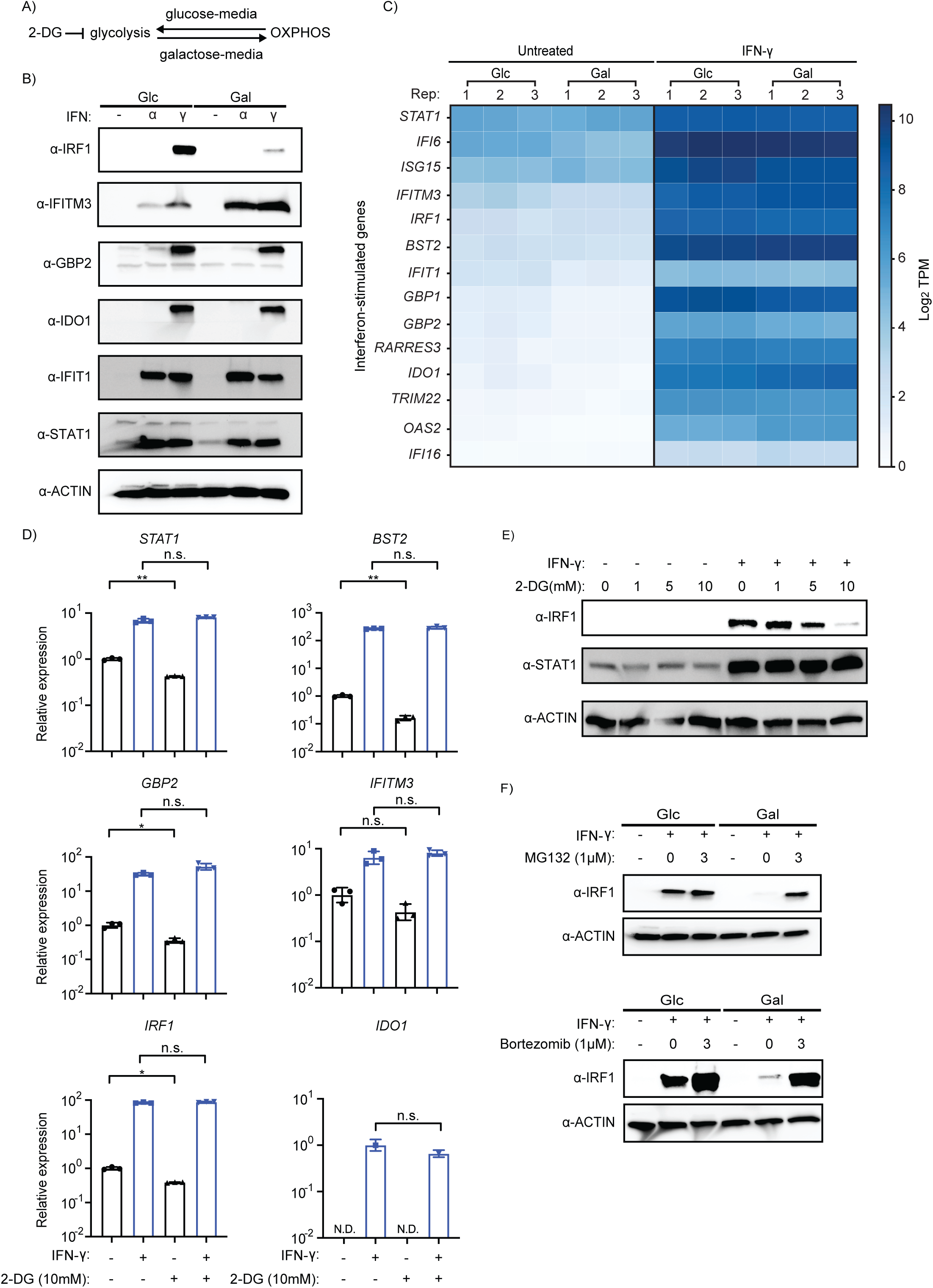
Glucose/galactose selectively alters ISG protein not RNA levels in interferon-primed cells. (A) Schematic of glucose/galactose culture conditions. 2-DG is a glycolytic inhibitor. (B) Western blot analysis of canonical ISGs in A549 cells treated with IFN-α or IFN-γ in either glucose (25 mM) or galactose (10 mM) media. Glc: glucose. Gal: galactose. Loading control for western blot: β-actin. Concentration for IFN-α and IFN-γ: 1000 units/mL. (C) RNA-seq analysis of canonical ISGs from cells pre-treated with IFN-γ, grown in glucose or galactose (N = 3 per treatment); log_2_TPM values. (D) qPCR of canonical ISGs relative to *β-actin* for IFN-γ-primed cells in glucose media treated with glycolysis inhibitor 2-DG, 10 mM (N = 3). (E) Western blot analysis of A549 cells in glucose media treated with various concentrations of 2-DG for 24 hours in the presence of IFN-γ (1000 U/mL). (F) Western blot analysis using lysates from A549 cells treated with the proteasome inhibitors - MG132 (top) and bortezomib (bottom) - and primed with IFN-γ in glucose/galactose. Either 1 μM MG132 or 1 μM bortezomib was added to the culture for 0 and 3 hours at 48 hours post-IFN-γ treatment. Statistical analysis was performed using an unpaired t-test in GraphPad Prism 9.5.1: n.s. not significant, * P ≤ 0.05, ** P < 0.01, *** P ≤ 0.001, **** P ≤ 0.0001.

To characterize the impact of glucose/galactose on type I (IFN-α) and type II (IFN-γ) ISG responses in A549 cells, we performed western blot on a subset of ISGs with known antiviral and immunomodulatory roles (Fig 3B). Both IFN-α and IFN-γ activities are known to be antagonized by diverse viruses commonly used as models that have biomedical relevance like the prototypical poxvirus, vaccinia [60,61]. A549 lung cells, which are derived from a barrier tissue, are a common model for diverse viruses including influenza A, coronaviruses, and vaccinia virus (VACV), as these cells have most host defense pathways intact [62]. We primed cells for twenty-four hours with interferon treatment followed by media replacement and harvested twenty-four hours thereafter. These conditions were selected to inform potential responses in bystander cells and to align with possible viral infection studies. Unexpectedly, marked differences were detected for two ISG proteins - IRF1 and IFITM3 – in cells with extended IFN stimulation grown in glucose relative to galactose. Interestingly, IRF1 is increased in glucose/IFN-γ lysates relative to galactose/IFN-γ (Fig 3B). Contrastingly, yet equally striking, IFITM3 (interferon-induced transmembrane protein 3) is enhanced in galactose media relative to glucose media in both IFN-α and IFN-γ treated cells (Fig 3B). The other ISG proteins tested - STAT1, GBP2, IFIT1, IDO1 - did not display similar differences in accumulation between glucose/galactose.

Intrigued by the differential expression of IRF1, which is known to inhibit diverse viruses [63], we next examined the impact of glucose/galactose on ISG RNA levels by performing RNA-seq on IFN-γ treated cells (Fig 3C). These data showed no major differences in RNA levels of ISGs including *IRF1* or *IFITM3*, which was validated by qPCR (S2B Fig). Altogether, these findings suggest that glucose/galactose alters the abundance of two IFN-induced ISGs largely at the protein, and not RNA, level at this time point.

To gain further insights into the interplay between glucose/galactose and interferons, we performed a time course analysis across conditions for IRF1 and IFITM3 RNA and protein (S3 Fig). This experiment showed differential expression kinetics for IRF1 and IFITM3 in IFN-α and IFN-γ treated cells. For instance, *IRF1* RNA is increased by both interferons as early as four hours but largely only IFN-γ at forty-eight hours (S3A Fig). IRF1 protein upregulation (four hours IFN-γ treatment) was detected earlier than IFITM3 (both IFN-α and IFN-γ). IRF1 protein was upregulated more strongly by IFN-γ than IFN-α across all time points. At earlier time points, IRF1 protein was detectable in galactose/IFN-γ but was noticeably decreased compared to glucose/IFN-γ at forty-eight hours (S3B Fig). Interestingly, elevated IFITM3 protein levels also visibly diverged in the media conditions by forty-eight hours post-treatment (S3B Fig). This experiment suggests interplay between interferons, duration of priming, and glucose/galactose conditions.

Given IFN-γ induced IRF1 is increased in conditions that favor glycolysis (glucose media) during extended IFN-γ priming, we next tested the impact of 2-deoxyglucose (2-DG) treatment. 2-DG is a glycolytic inhibitor that is known to block hexokinase and often used to complement glucose/galactose studies (Fig 3A). 2-DG treatment (10 mM) of glucose/IFN-γ treated cells resulted in an almost complete loss of IRF1 protein but not *IRF1* RNA expression (Fig 3D, Fig 3E). The same treatment had no noticeable effect on induced *STAT1* RNA or STAT1 protein levels. In addition, qPCR of four other ISGs showed no major effects of 2-DG/IFN-γ treatment on these transcripts (Fig 3D). These data further support that glucose influences IFN-γ induced IRF1 protein levels.

To distinguish whether the differential regulation of IFN-γ induced IRF1 is due to alterations in protein translation or turnover, we treated A549 cells with two different types of proteasome inhibitors - MG132 and bortezomib. Both MG132 and bortezomib treatment (3 hours) dramatically increased IRF1 levels in galactose/IFN-γ primed cells (Fig 3F). We excluded a role for global protein turnover in the observed changes in IRF1 associated with glucose/galactose as no differences were observed under the same conditions for a set of proteins that localize to distinct cellular compartments - ACTIN (cytosol), HDAC1 (nuclear), SDHA (mitochondrial inner membrane), and TOM70 (mitochondrial outer membrane) (S2C Fig). These data suggest that IRF1 undergoes selective proteasomal degradation in response to extended galactose/IFN-γ culture.

### Regulation of IFN-γ induced IRF1 protein levels by glucose/galactose is conserved in mammals

Evolutionary signatures inform the relevance of biological phenomena. IRF1 is highly conserved as evidenced by homologs in oysters [64]. To test whether differential regulation of IRF1 by glucose/galactose is conserved in mammals, we analyzed *IRF1* RNA and protein levels in primary mouse embryonic fibroblasts (MEFs; E15) and feline kidney cells (CRFK) grown in either condition followed by interferon priming. For both cell types, we saw robust induction of *IRF1* RNA by IFN-γ in both glucose and galactose conditions (Fig 4A, Fig 4B). Notably, we saw increases comparable to A549 cells for IFN-γ induced IRF1 protein for both mouse (Fig 4C) and cat cells (Fig 4D) in glucose media relative to galactose media. No similar trend for cat and mouse cells was observed for other tested ISGs like STAT1. These data indicate that differential regulation of IFN-γ induced IRF1 protein by glucose/galactose is conserved in unrelated cell types derived from animals separated by nearly one hundred million years of evolutionary divergence.

**Fig 4.**
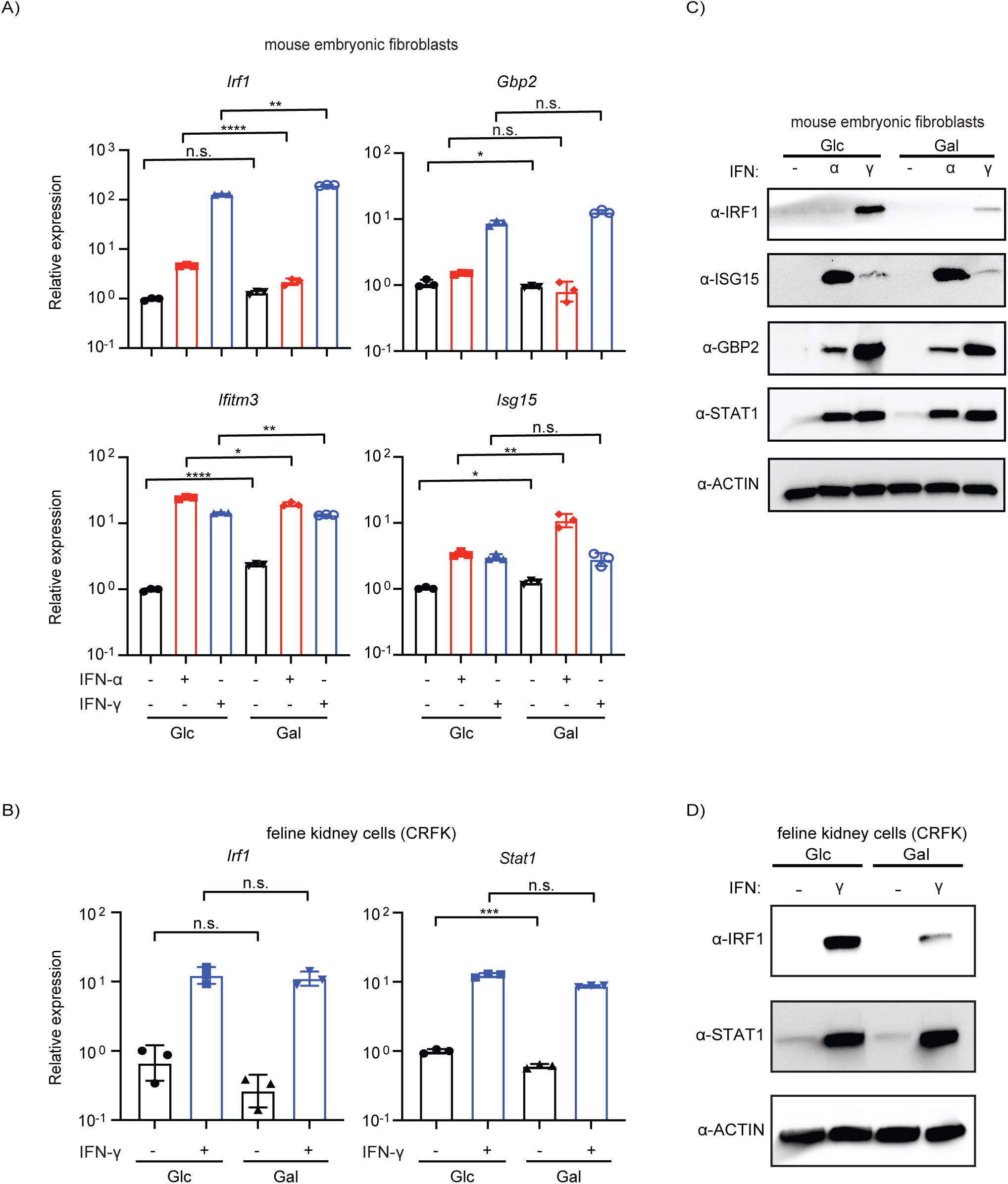
Glucose/galactose regulation of IFN-γ induced IRF1 protein is conserved in mammals. (A) qPCR analysis of ISGs from mouse embryonic fibroblasts (MEFs, E15) pre-treated with IFN-α or IFN-γ grown in media containing either glucose or galactose. Concentration for IFN-α and IFN-γ: 1000 units/mL. (B) qPCR analysis of *IRF1* and *STAT1* from IFN-γ-primed Crandell-Rees Feline Kidney (CRFK) cells grown in either glucose or galactose. Concentration for IFN-γ: 1000 units/mL. (C) Western blot analysis of ISGs for MEFs primed with IFN-α or IFN-γ in glucose or galactose media. Concentration for IFN-α and IFN-γ: 1000 units/mL. (D) Western blot analysis of IRF1 and STAT1 for CRFK cells pre-treated with IFN-γ in glucose or galactose media. Concentration for IFN-γ: 1000 units/mL. Glc: glucose (25 mM). Gal: galactose (10 mM). Loading control for western blot: β-actin. Statistical analysis was performed using an unpaired t-test in GraphPad Prism 9.5.1: n.s. not significant, * P ≤ 0.05, ** P < 0.01, *** P ≤ 0.001, **** P ≤ 0.0001.

### Glucose/IFN-γ, but not galactose/IFN-γ, inhibits vaccinia virus replication

To examine functional consequences of glucose/galactose and differential regulation of IFN-γ induced IRF1, we infected human A549 lung cells that were primed with either IFN-α (type I interferon) or IFN-γ (type II interferon) with the prototypical poxvirus - vaccinia virus (timeline: S4A Fig). VACV is frequently used to study host defenses, in part, because it encodes numerous immunomodulators [65,66] including type I and type II interferon antagonists such as soluble receptor decoys for both IFN-α and IFN-γ [60]. While VACV and other poxviruses are highly successful in their ability to evade innate immunity, IFN-γ pretreatment can attenuate VACV replication in some cell types [67,68]. IRF1 is also a known restriction factor for VACV in certain settings [68,69].

First, we started by performing experiments using a VACV reporter strain [70] with a low but common MOI (MOI = 0.01) given vaccinia is known to extensively counter host defenses in culture. These experiments showed a significant reduction in expression of both an early (SSB) and late viral protein (A27) in glucose/IFN-γ relative to the other treatments including IFN-α primed cells (Fig 5A). Consistently, glucose/IFN-γ priming also resulted in a drastic reduction of other markers of viral replication. Specifically, we observed 1) decreased levels of VACV RNA [qPCR: early gene (*I3L*), late gene (*F17R*)] (Fig 5B), 2) decreased levels of vaccinia luciferase reporter expression (S4B Fig)], and 3) a ∼2 log reduction in infectious vaccinia titers relative to glucose untreated (Fig 5C). The continued, marked reduction in VACV replication at forty-eight hours post infection in IFN-γ primed, glucose-grown cells indicated a long-lasting activity specific to these conditions (S4B Fig). Reduced VACV replication was also observed when IFN-γ (S4C Fig) or glucose levels (S4D Fig) were decreased.

**Fig 5.**
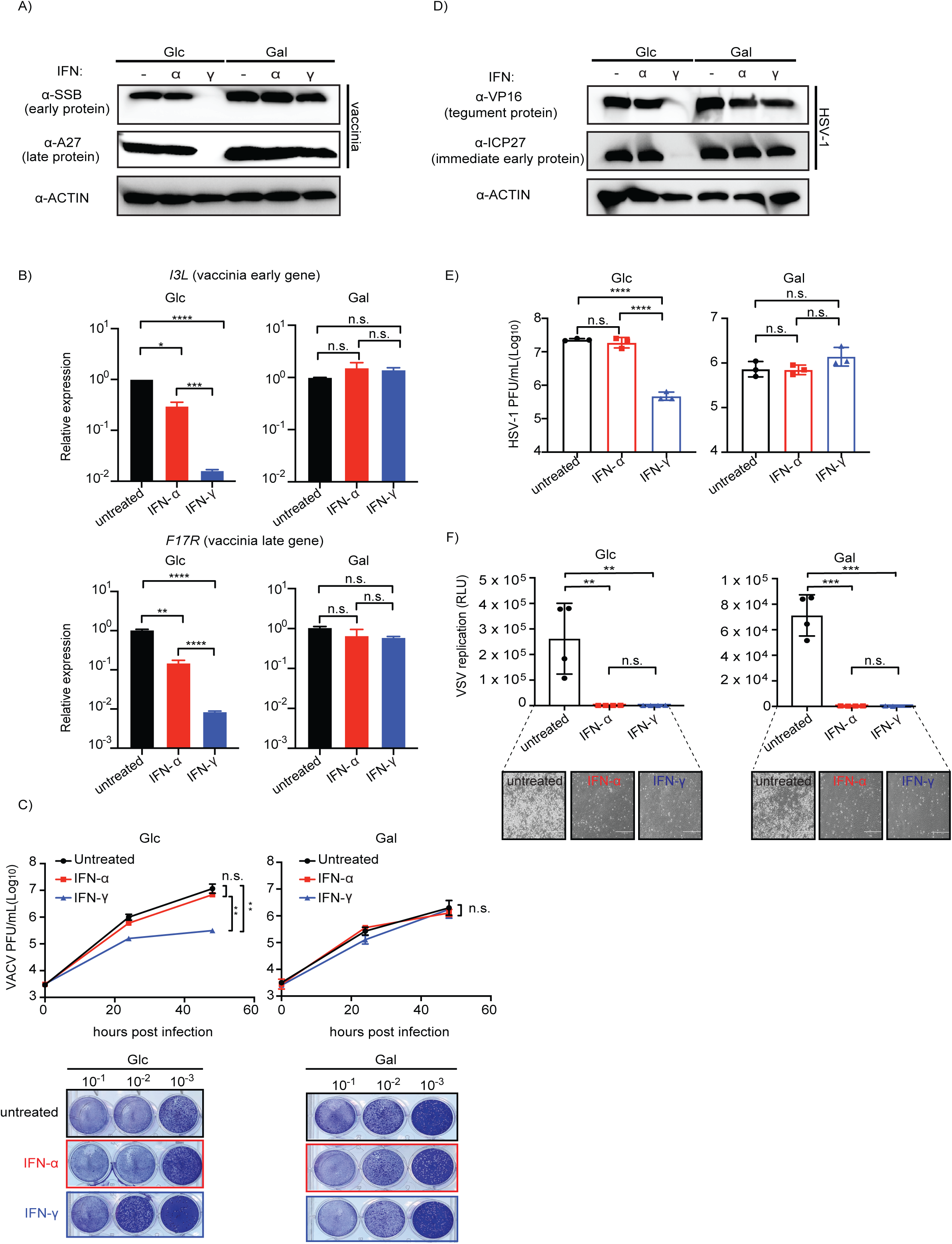
Glucose but not galactose culturing of IFN-γ primed cells reduces VACV and HSV-1 replication. (A) Western blot analysis of VACV proteins – early (SSB) and late (A27) – in A549 cells primed with either IFN-α or IFN-γ, grown in glucose or galactose. (B) qPCR of VACV transcripts – early (*I3L*) and late (*F17R*) genes (N = 3). (C) Top: Quantification of plaque assay for VACV-Luc-GFP infectious units from A549 cells (N = 3). Bottom: representative image of plaque assay with various dilutions of viral titers (10^-1^-10^-3^). (D) Western blot analysis for HSV-1 viral immediate early (ICP27) and tegument proteins (VP16) in A549 cells. (E) Quantification of plaque forming units from HSV-1-GFP A549 infected cells primed with either glucose or galactose as well as IFN-α or IFN-γ (N=3). (F) VSV (luciferase) replication assay from IFN-pretreated A549 cells grown in glucose or galactose. Bottom: images showing presence or absence of cytopathic effect (with interferon priming). Plaque forming unit: P.F.U. Glc: glucose (25 mM). Gal: galactose (10 mM). Loading control for western blot and qPCR: β-actin. For viral infection, MOI = 0.01 for VACV-Luc-GFP and VSV-Luc; MOI = 1 for HSV-1-GFP. Concentration for IFN-α and IFN-γ: 1000 units/mL. Statistical analysis was performed using an unpaired t-test in GraphPad Prism 9.5.1: n.s. not significant, * P ≤ 0.05, ** P < 0.01, *** P ≤ 0.001, **** P ≤ 0.0001.

To gain increased resolution regarding what vaccinia lifecycle stage is affected by the glucose/IFN-γ response, we performed experiments using a high vaccinia MOI (MOI = 3) (S5A Fig, S5B Fig). With a high MOI, we observed increased vaccinia reporter expression in glucose/IFN-γ (S5A Fig) comparable to that of glucose alone and glucose/IFN-α. The increased reporter expression, which is driven by a synthetic poxvirus p7.5 early/late promoter, was corroborated by increased levels of the vaccinia early protein SSB in glucose/IFN-γ (S5B Fig). In contrast, levels of the vaccinia late protein A27 were still noticeably decreased similar to MOI = 0.01 in glucose/IFN-γ relative to glucose alone and glucose/IFN-α. Thus, at low MOIs the response in glucose/IFN-γ reduced markers of early and late viral replication whereas at a higher MOI the conditions still limited vaccinia late viral protein expression. Thus, vaccinia virus replication is reduced under conditions that mirror expression of IRF1; namely glucose/IFN-γ.

The plaque assay data highlighted that there was also an overall one-log decrease in vaccinia titers in galactose media relative to glucose media. A similar trend of reduced vaccinia replication in galactose, albeit to a lesser extent, was evident for the reporter virus assays (S4B Fig, S5A Fig). These data combined with comparable expression of early and late vaccinia proteins across glucose/galactose (Fig 5A, S5A Fig), excluding glucose/IFN-γ, suggest the presence of a disruptive activity or deficiency at a late stage in the viral lifecycle associated with galactose.

To test the generality of our findings, we performed infections using similar conditions with the unrelated herpes simplex virus-1 (HSV-1; MOI = 1). HSV-1 replicates in the nucleus in contrast to the cytoplasmically replicating vaccinia [71]. Like vaccinia in glucose/IFN-γ conditions, HSV-1 displayed marked decreases in viral protein production and a two-log decrease in infectious titers relative to glucose untreated (Fig 5D, Fig 5E). The near loss of expression for immediate early protein ICP27 in glucose/IFN-γ relative to the other treatments suggests HSV-1 replication is blocked at an early stage of HSV-1 infection (Fig 5D). Similar to vaccinia, galactose supplementation led to decreased HSV-1 viral titers (∼one log by plaque assays, Fig 5E, S4E Fig) but not lower HSV-1 protein expression, suggesting a disruptive activity or deficiency at a late stage in HSV-1 replication. To further explore glucose/galactose activity on viral replication, we performed infections with the model RNA virus, vesicular stomatitis virus (VSV; MOI = 0.01). These infections, in contrast, showed that glucose/galactose did not perturb the ability of either IFN to potently inhibit VSV replication. Specifically, we observed both a loss of VSV luciferase reporter expression and cytopathic effect (Fig 5F) in cells primed with either IFN-α or IFN-γ irrespective of glucose/galactose.

### Glucose/galactose does not result in overt changes in cell viability

To exclude that the reductions in viral replication associated with glucose/IFN-γ and across galactose conditions were due to differences in cellular viability, we interrogated markers of cell death. First, we imaged uninfected and vaccinia infected cells across our test conditions: glucose/galactose with or without IFN-priming (Fig 6A, Fig 6B). These images displayed no major differences in monolayer formation or dying (floating) cells for uninfected cells in either glucose/galactose with or without IFN-priming. For vaccinia infected cells, these images showed visible cytopathic effect for all glucose and galactose conditions except for glucose/IFN-γ cells which still appeared largely as an intact monolayer (Fig 6A, Fig 6B). Next, we analyzed levels of cleaved PARP by western blot in uninfected (Fig 6C) and vaccinia infected cells (Fig 6D). In uninfected cells, minimal but variable levels of PARP cleavage across conditions were observed. In vaccinia infected cells, the largest increase in PARP cleavage relative to the other conditions was associated with glucose/IFN-γ (Fig 6D). The increased PARP cleavage could be due to enhanced cell death or reduced vaccinia antagonism of pathways that regulate PARP cleavage because of decreased viral replication. To account for potential differences in the efficiency of PARP cleavage between glucose/galactose, we infected cells with VSV. This experiment demonstrated that glucose/galactose did not noticeably impact the dramatic levels of PARP cleavage induced by VSV infection (Fig 6D). Collectively, these data suggest that 1) an activity is present in glucose/IFN-γ, but not galactose/IFN-γ, which attenuates VACV replication at time point that influences late protein expression even at high MOIs and 2) that galactose relative to glucose culture results in decreased production of infectious vaccinia and HSV-1 at a lifecycle stage post-gene expression.

**Fig 6.**
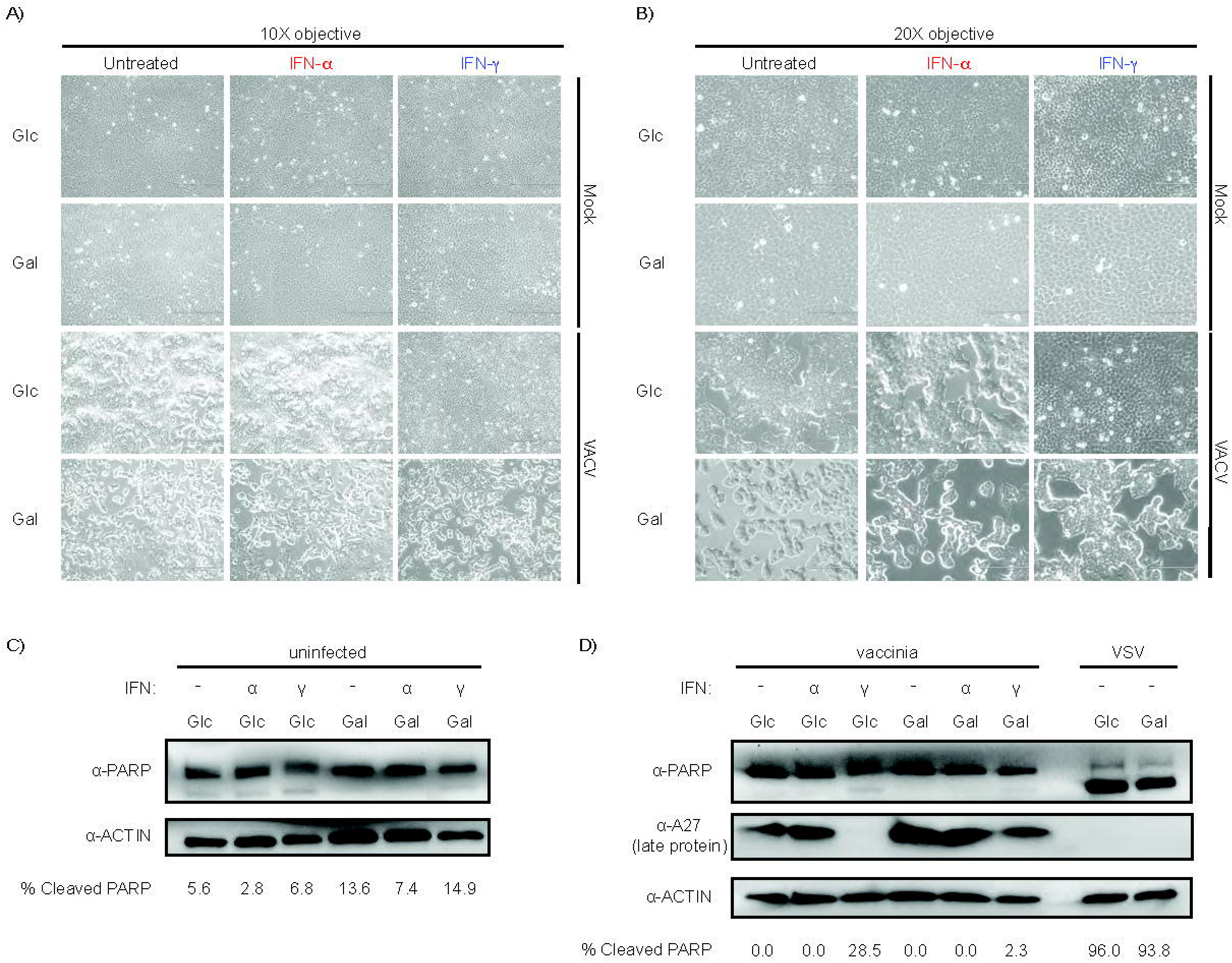
Reductions in vaccinia virus replication do not correlate with markers of cell viability. (A) Brightfield microscope images at 10X of A549 cells grown in glucose/galactose with and without interferon priming; uninfected (top) and vaccinia virus infected (bottom). (B) Brightfield microscope images at 20X of A549 cells grown in glucose/galactose with and without interferon priming; uninfected (top) and vaccinia virus infected (bottom). (C) PARP cleavage western blots of uninfected A549 cells grown in glucose/galactose with and without interferon priming. (D) PARP cleavage western blots of vaccinia virus infected A549 cells grown in glucose/galactose with and without interferon priming. Control: VSV-infected glucose/galactose cells (no interferon priming). Glc: glucose (25 mM). Gal: galactose (10 mM). Loading control: β-actin. A27: vaccinia virus late protein. IFN-α or IFN-γ concentration: 1000 units/mL.

### *IRF1* impacts VACV replication in glucose/IFN-γ but not across galactose conditions

In glucose/IFN-γ, possible explanations for the decreased vaccinia replication include that glucose, but not galactose, potentiates IFN-γ - instead of IFN-α - signaling. Two ways to modulate the antiviral effect of IFN-γ are by 1) increasing the overall magnitude of the IFN-γ induced response or 2) altering the composition of the antiviral response at the RNA or protein level. To distinguish between these possibilities, we first assayed steady-state levels of total STAT1 and activated STAT1 (phosphorylated Tyr701) by western blot across conditions. This experiment revealed no marked differences for either total levels of STAT1 or phospho-STAT1 (Tyr701) in uninfected or VACV-infected cells grown in either media with or without IFN-γ priming (S6A Fig). Next, we examined changes in gene expression mediated by glucose/galactose in IFN-γ primed cells.

Analysis of our RNA-seq dataset matching the media and treatment conditions herein, we identified 312 genes upregulated common to both glucose/IFN-γ/mock infected and glucose/IFN-γ/vaccinia infected conditions (S6B Fig, S2 and S3 Tables). We also identified 176 genes upregulated in both galactose/IFN-γ/mock infected and galactose/IFN-γ/vaccinia infected conditions. GO analysis of the differentially expressed genes did not identify signatures indicative of interferon and/or interferon responses (S6C Fig). The top three pathways enriched for genes upregulated in glucose/IFN-γ conditions (S6C Fig) were extrinsic prothrombin activation pathway, coagulation system, and pulmonary fibrosis idiopathic signaling pathway. As the GO categories did not point to any appreciated regulators of the host response, we next considered the possibility of atypical antiviral factors contributing to the observed glucose/IFN-γ response. To do so, we selected four factors for follow-up from our list of differentially expressed genes that displayed log_2_ fold change > 2 in the overlap of mock and vaccinia infected glucose/IFN-γ relative to overlap of mock and vaccinia infected galactose/IFN-γ conditions (adjusted p-value ≤0.01) (S6B Fig): *TXNIP*, *PLA2G2A*, *SCN4A*, *SUCNR1*. We validated the changes in RNA-seq of these four factors by qPCR (S6D Fig). However, these RNA changes did not result in similar changes in protein levels for these factors. Specifically, protein levels of TXNIP and SUCNR1, which were assayed using commercially available antibodies, were discordant with the antiviral activity associated with glucose/IFN-γ (S6E Fig).

Given the correlation of IRF-1 expression and the response in glucose/IFN-γ, we next considered the role of IRF1 in contributing to the differences in infection outcomes associated with glucose/galactose. Overexpression of IRF1 results in the inhibition of diverse viruses [63] including vaccinia [68,69]. In addition to known roles in amplification of the ISG response by direct DNA-binding [63,72], IRF1 activity is associated with expression of inflammatory genes [73], DNA-repair genes [74] as well as gene induction by a scaffolding mechanism independent of DNA-binding [75].

To interrogate the impact of IRF1 on vaccinia replication in glucose/galactose conditions, we generated polyclonal *IRF1* knockout (KO) A549 cell lines using CRISPR/Cas9 delivered by lentivirus (Fig 7A). As controls, polyclonal *STAT1* KO A549 cell lines were also generated (S7A Fig). VSV infections were used as a control given that glucose/galactose did not influence infection outcomes under the conditions tested (Fig 5F). In *STAT1* KO cells, IFN-γ treatment showed a reduced effect on blocking VSV (S7B Fig) and VACV (S7C Fig) replication. These data are consistent with an IFN-γ induced STAT1-mediated response limiting vaccinia replication in glucose. In *IRF1* KO cells, no increases in VSV replication were observed in glucose/IFN-γ (Fig 7B) or galactose/IFN-γ cells (Fig 7C). These results are in agreement with previous findings showing that multiple ISGs are sufficient to block VSV [76] and our data above showing that glucose/galactose did not display detectable effects on VSV infection under the conditions tested (Fig 5F).

**Fig 7.**
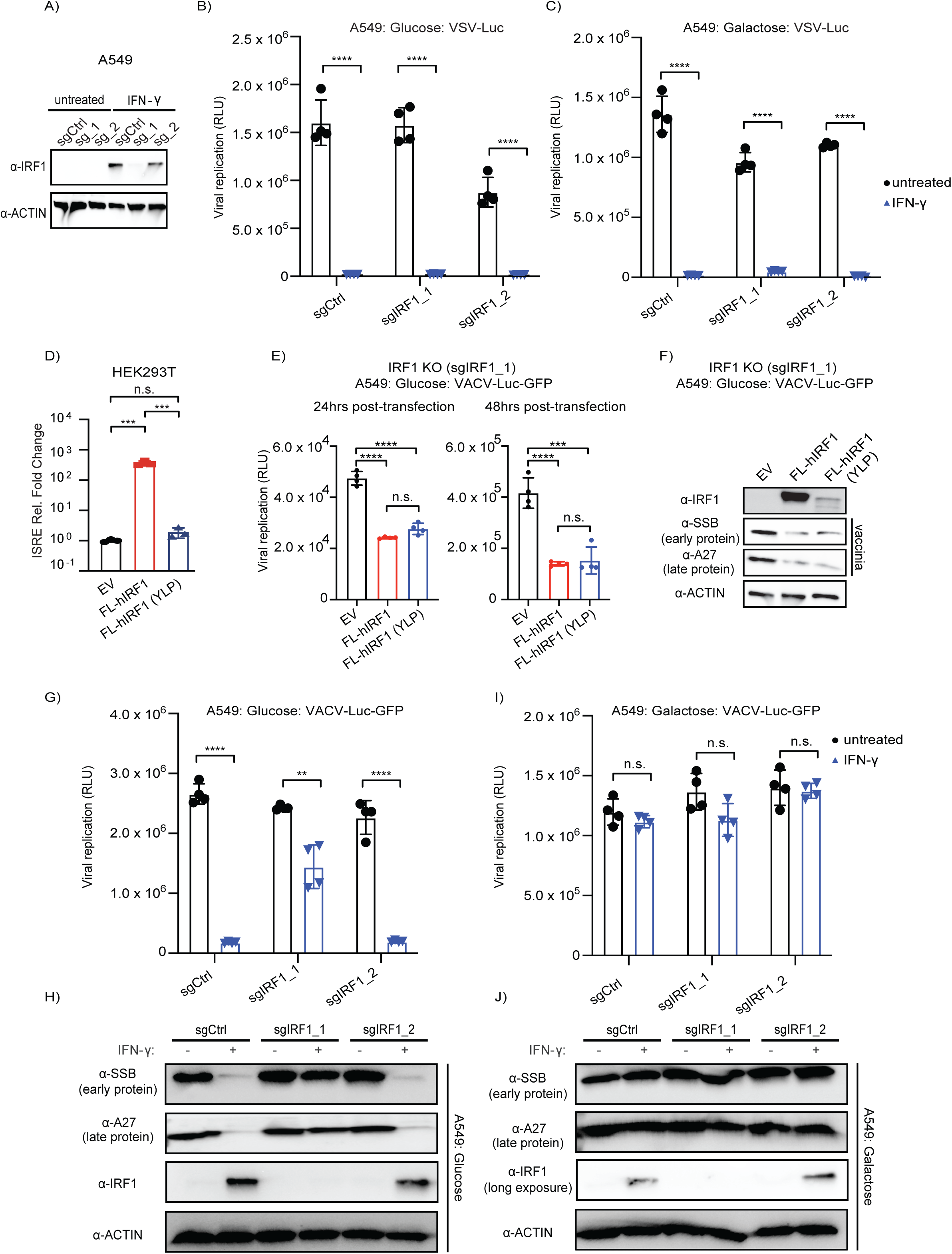
*IRF1* impacts VACV replication in glucose/IFN-γ primed but not galactose conditions. (A) Western blot analysis of *IRF1* KO A549 cell lines generated with pLentiCRISPR V2 vectors. Two sgRNAs were used per target gene; sgCtrl – non targeting control. (B) Glucose: VSV-Luc (luciferase) replication assays in IRF1 *KO* cells primed with IFN-γ (N = 4). (C) Galactose: VSV-Luc (luciferase) replication assays in *IRF1 KO* cells primed with IFN-γ (N = 4). (D) ISRE reporter assay of 293T cells transfected with human FLAG-IRF1 plasmids. EV: empty vector, FL-hIRF1: WT human FLAG-IRF1, FL-hIRF (YLP): human FL-IRF1 Y109A/L112A/P113A mutant [75]. (E) Glucose: no IFN-priming vaccinia (luciferase) replication assays - 24 hours post infection - in *IRF1* KO cells (sgIRF1_1) transiently transfected with human FLAG-IRF1 plasmids for 24 hours or 48 hours prior to infection (N = 4); VACV-Luc-GFP (MOI = 0.01). EV: empty vector, FL-hIRF1: WT human FLAG-IRF1, FL-hIRF1 (YLP): human FL-IRF1 Y109A/L112A/P113A mutant [75]. (F) Western blot analysis of glucose: no IFN-priming VACV-infected *IRF1* KO cells transiently transfected with human FLAG-IRF1 plasmids. Infection was performed with VACV-Luc-GFP (MOI = 0.01) at 48 hrs post-transfection, and protein was harvested at 24 hours post-infection. (G) Glucose: VACV-Luc-GFP (luciferase) replication assays in *IRF1 KO* cells primed with IFN-γ (N = 4). (H) Glucose: western blot of vaccinia virus infected *IRF1* KO A549 cells with or without IFN-γ priming. (I) Galactose: VACV-Luc-GFP (luciferase) replication assays in *IRF1 KO* cells primed with IFN-γ (N = 4). (J) Galactose: western blot of vaccinia virus infected *IRF1* KO A549 cells with or without IFN-γ priming. IRF1 blot in galactose is long exposure. Statistical analysis was performed using an unpaired t-test in GraphPad Prism 9.5.1: n.s. not significant, * P ≤ 0.05, ** P< 0.01, *** P ≤ 0.001, **** P ≤ 0.0001.

To test the genetic contribution of *IRF1* to the regulation of vaccinia replication across glucose/galactose conditions, we analyzed vaccinia replication in WT and *IRF1* KO cells in a series of experiments. First and as a control, we transiently transfected a recoded WT IRF1 and the IRF1 DNA-binding mutant (Y109A/L112A/P113A; YLP); both with N-terminal FLAG tags. IRF1 (YLP) is reported to be deficient in ISRE-binding but capable of activating gene expression by a putative scaffolding mechanism [75]. Consistently, we confirmed WT IRF1, but not IRF1 (YLP), is able to activate an ISRE-luciferase reporter (Fig 7D). In sgIRF1_1 KO cells in glucose/no interferon-priming, both WT IRF1 as well as IRF1 (YLP) reduced vaccinia reporter (Fig 7E) and vaccinia virus protein expression (Fig 7F). Under other conditions transient transfection and stable IRF1 expressing cells resulted in cellular toxicity in WT and *IRF1* KO cells.

To examine vaccinia replication and its effect on host responses in *IRF1* KO cells treated with IFN-γ compared to control, we analyzed levels of cleaved PARP, took images of uninfected and infected cells, and assayed markers of vaccinia replication. Western blot analysis indicated cleaved PARP was elevated in both sgCtrl and sgIRF1_1 KO uninfected and vaccinia infected cells in glucose and galactose when primed with IFN-γ (S8 Fig). These cleaved PARP patterns in IFN-γ treated cells were largely abrogated in *IRF1* KO cells (S8B Fig, S8D Fig). Consistent with an increase in vaccinia replication, images of infected sgIRF1_1 KO cells in glucose/IFN-γ displayed increased cytopathic effect relative to sgCtrl cells, (S9 Fig). In comparison, sgIRF1_1 KO cells in galactose displayed similar cytopathic effect irrespective of IRF1 genotype and interferon-priming. These data suggest that IRF1 is implicated in elevated PARP cleavage in IFN-γ treated cells, which may be due, in part, to an aspect of CRISPR/Cas engineering with lentivirus. In addition, there also appears an association between cytopathic effect and IRF1 specific to glucose/IFN-γ, which differs from the PARP cleavage pattern that occurs in sgCtrl treated IFN-γ cells in glucose and galactose.

Next, we tested the impact of IRF1 loss on vaccinia virus replication. We found that vaccinia virus showed increased replication in sgIRF1_1 KO cells relative to sgCtrl in glucose/IFN-γ conditions by viral reporter (Fig 7G) and western blot for viral proteins (Fig 7H). As a control, we also infected cells where *IRF1* knockout was inefficient (sgIRF1_2 KO); these cells displayed negligible changes in VACV replication in glucose/IFN-γ relative to sgCtrl group. Likewise, we also observed increased expression of vaccinia RNAs in glucose/IFN-γ for sgIRF1_1 KO cells but not WT or sgIRF1_2 KO cells (S10 Fig). In contrast to glucose conditions, vaccinia replicated to comparable levels in sgCtrl, sgIRF1_1 KO, and sgIRF1_2 KO cells in galactose conditions with or without IFN-γ priming as evidenced by data for 1) vaccinia reporter virus assays (Fig 7I), 2) the early protein, SSB, and 3) the late protein, A27 (Fig 7J). Collectively, these data indicate that the genetic status of *IRF1* tips vaccinia replication in glucose media where IFN-γ induced IRF1 protein levels are high but not galactose conditions where IFN-γ induced IRF1 levels are markedly reduced.

## DISCUSSION

### Human tonic ISG expression displays subclass substructure across tissues

Our study is rooted in the characterization of ISG responses and factors that influence them. In the first part of the study, we focused on tonic ISG RNA expression across human tissues. To examine factors that may contribute to differences in ISG expression like metabolism, which varies across tissues and cell-types, we developed a cell culture protocol adapted from the mitochondrial field. Unexpectedly, we found that differential stabilization of interferon-γ induced IRF1 protein by glucose/galactose culture conditions (Fig 3) is associated with distinct vaccinia infection outcomes in those same conditions (Fig 5, Fig 7). Often viewed as a single class of genes, ISGs are appreciated for their roles in host defense and shaping susceptibility to pathogen infection [17,22]. Expression of canonical ISGs is also commonly leveraged as a signature of innate immune activation in various contexts including human disease [25]. Still, the severity of viral disease [29,77] seems to be increasingly linked to variability in the repertoires of expressed ISGs as evidenced by isoforms of OAS1 as well as OAS1 loss-of function alleles associated with distinct outcomes in COVID-19 patients. However, our understanding of the factors, including non-genetic cues, contributing to the composition of the ISG response remains limited in comparison to the factors shaping the magnitude of the response. To explore unappreciated layers of ISG regulation, a more cohesive snapshot of ISG expression has been needed.

Although variation in induced ISGs has been described for a subset of cell types, tissues, and species [19,34,39,78,79], data for ISG expression across an organism, which includes barrier tissues, has been lacking. To this end, we analyzed tonic ISG expression (Fig 1, Fig 2) as a proxy to inform and generate hypotheses related to regulatory mechanisms of induced ISGs. We report that tonic ISG expression is highly variable and seemingly independent of interferon and interferon receptor RNA levels across human tissues (Fig 1). This finding contributes to an emerging view that constitutive and robust expression of ISGs is essential for host defense [34,35,80]. This analysis highlights an uncharacterized division among human (tonic) ISGs - high, low, and variable subclasses (Fig 1). These subclasses foreshadow ill-defined regulatory mechanisms for a response considered a hallmark of many genetic and complex diseases [25].

### Glucose/galactose influences stability of the key ISG protein IRF1

By applying a common methodology leveraged in mitochondrial research as a tool to study the impact of metabolic rewiring on host defenses (Fig 3A), we uncovered an unappreciated and conserved layer of regulation for IRF1 - a key ISG protein. The functional relevance of glucose/galactose regulation, and seemingly glycolysis (Fig 3D, Fig 3E), of IFN-γ induced IRF1 protein is supported by the differences in vaccinia replication in *IRF1* KO cells between the two conditions (Fig 7). While IRF1 antiviral activity is well-demonstrated for VACV and other viruses, it has largely been presumed that this occurs through the transcriptional activation of ISRE-containing promoters but to our knowledge this has not been formerly tested. Interestingly, overexpression of WT and IRF1 YLP mutant in *IRF1* KO cells results in similar decreases in vaccinia replication which suggests that IRF1 ISRE binding activity may be dispensable at least during vaccinia infection (Fig 7E, Fig 7F). This finding may partially account for the lack of both “usual (ISG) suspects” and differentially expressed genes with IRF1-binding sites (S11 Fig, S4 Table) in the RNA-seq (Fig 3C), while implying that IRF1 may regulate at least some viruses by other less defined activities.

One possibility is that IRF1 acts by modulating cell death - which is a major host defense mechanism [55,81] that can also be proviral [82], - via interplay between interferons and metabolism. Our data indicate that the elevated PARP cleavage in both glucose/IFN-γ and galactose/IFN-γ sgCtrl cells when uninfected and vaccinia-infected is likely linked to IRF1 (S8 Fig). However, the cleaved PARP pattern appears potentially distinct from the reduced cytopathic effect (S9 Fig) and decreased vaccinia replication associated with IRF1; both of which are exclusive to glucose/IFN-γ. Future work will potentially inform IRF1 (YLP) activities that shape vaccinia infection outcomes.

Our studies also provide evidence that IFN-α and IFN-γ induced IFITM3 protein is also regulated by glucose/galactose (Fig 3B, Fig 3C, S3 Fig). The directionality by which IRF1 and IFITM3 are regulated by glucose/galactose is intriguing in light of the functions associated with these two ISGs. IRF1 is an activator of the inflammatory response whereas IFITM3 is an anti-influenza A virus [83,84] and anti-SARS-CoV2 factor [85]. Our data may have relevance for certain metabolic disorders and mitochondrial diseases which are often characterized by mitochondrial dysfunction as well as altered glucose metabolism. Individuals with these conditions commonly display exacerbated inflammation during viral infection and increased susceptibility to certain viruses including influenza A virus and SARS-CoV2 [86–88]. Future studies including *in vivo* studies will increase our understanding of post-translational regulation of ISG proteins by glucose/galactose and potentially other metabolites. Distinctly, our experimental protocol may be useful in developing future genetic screens to uncover additional host defense mechanisms as many ISGs remain uncharacterized.

Degradation of select ISGs is a key strategy used by a range of viruses like HIV-1 and poxviruses to suppress interferon-stimulated host defenses [89–91]. Unexpectedly, our studies illustrate that non-viral cues, in this case glucose/galactose, can influence the composition of ISG responses and result in distinct infection outcomes. This finding contrasts with an “all-hands-on-deck” strategy deploying all ISGs during infection. In addition to the protein level regulation of ISG proteins we report here, future studies might also consider the impact of feedback loops triggered by interferon treatment resulting in changes in metabolic flux - both in utilization and output – on the composition of ISG responses. Collectively, this study provides evidence for an unappreciated and conserved layer of regulation for a critical ISG protein which may have implications for the interpretation of ISG transcriptional signatures.

### Glucose/galactose as secondary cues that regulate the adaptability of ISG responses

Glucose and galactose were used to model secondary cue activities in interferon-primed infected cells because they are established tools to perturb mitochondrial functions [56,57,92]. Future experiments will distinguish whether glucose and galactose, their derivatives, or activities associated with these molecules serve as the actual triggers. The potency of these cues linked to mitochondrial activities is enticing due to documented crosstalk between host defenses and this organelle [54,55,93–100]. Indeed, there is a growing number of instances where metabolites and modulators of metabolism regulate transduction of immune signals to influence the magnitude of the response [31,101] and infection outcomes [102,103].

Lastly, there has been an emerging picture that metabolism impacts replication of viruses including vaccinia [44,55]. Many of these studies provide evidence indicating that vaccinia is actively promoting specific metabolic states and/or counteracting metabolic reprogramming. For instance, ribosomal profiling of vaccinia-infected cells has shown an increase in the translation of mRNAs encoding OXPHOS factors [104]. Several studies describe roles for changes in specific metabolites and metabolic flux during vaccinia replication including 1) levels of the TCA cycle metabolite citrate mediated by vaccinia encoded growth factor [105], 2) fatty oxidation [106], 3) asparagine [107], and glutamine [108]. More recently, vaccinia has been shown to target and modulate activities of the nutrient sensor, mTOR, via vaccinia-encoded F17 to influence ISG responses and glycolysis [109–111]. Given the connectivity of metabolic circuitry, it is likely that there is overlap between our findings with ISG responses and these previous reports. Perhaps certain virus-mediated changes in metabolism during infection occur, in part, as a means to counteract dynamic ISG responses mediated by the cell during infection. It is tempting to speculate that regulation of certain proteins by specific metabolite levels, which may change as a consequence of pathogen replication and inflammation, may represent a rapid means to invoke antiviral subprograms to adapt to an evolving infection (Fig 8).

**Fig 8.**
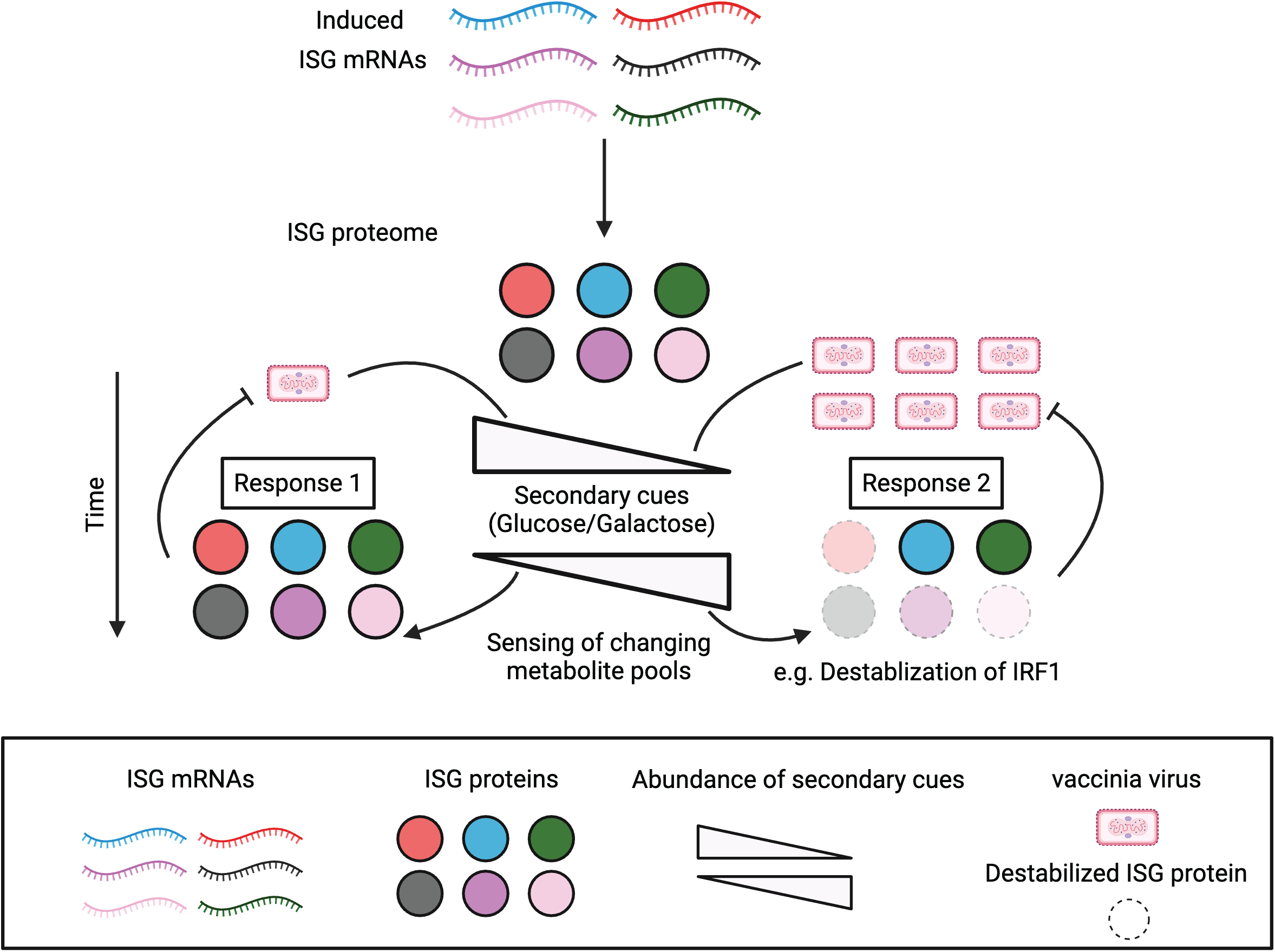
Proposed model for adapting host responses over time via changes in levels of select ISG proteins regulated by secondary cues. Binding of IFN to its cognate receptor induces the transcription and translation of ISG mRNAs. The presence of secondary cues (glucose or galactose) can alter antiviral responses via selective stabilization of some but not all ISG proteins to shape ISG repertoires. Sensing of metabolite consumption during viral replication could result in a feedforward loop adapting ISG responses at the protein level to promote adaptation of antiviral defenses. Response 1: If the virus is replicating at low levels (left), it is consuming relevant metabolites at a reduced rate. Changes in cellular metabolite pools are sensed by an unknown cellular factor/s which trigger a response that impacts protein levels of some but not all immune defense proteins to manage the infection. Response 2: If the virus is replicating at high levels (right), the virus is consuming necessary metabolites at a rapid rate which leads to major changes in the concentrations of these metabolite pools. Sensing of these changes in metabolite pools results in adaptation of antiviral defenses by altering the protein composition of the antiviral response to counter increased viral replication which may involve increased recruitment of different immune cell types. Changes at the protein level may allow for rapid adaptation of transcriptionally-induced antiviral responses. The diagram was generated with BioRender.

## MATERIALS AND METHODS

### Cell culture and treatments

Cell lines used in these studies were: A549 cells (a generous gift from Dr. Susan Weiss, University of Pennsylvania), BSC40 cells (Dr. Don Gammon, UT Southwestern), Vero cells (ATCC), HEK 293T (ATCC), CRFK cells (ATCC). All cells were maintained at 37°C in a humidified incubator at 5% CO_2_. HEK293T and A549 cells were grown in DMEM high glucose with 1 mM sodium pyruvate [25 mM (Corning)]. CRFK cells were maintained in EMEM (Corning) with 1 mM sodium pyruvate [25 mM (Corning)]. Media were supplemented with FBS [10%, (Gibco)], L-glutamine [2 mM (Corning)], 1X Antibiotic-Antimycotic [Gibco], and 10 mM HEPES [Corning]. On the day of the experiment, both CRFK and A549 cells were grown in DMEM, no glucose, without sodium pyruvate (ThermoFisher) supplemented with either glucose (25 mM) or galactose (10 mM) – a standard concentration to enhance cellular OXPHOS [56,57]. E15 mouse embryonic fibroblasts (MEFs) were derived from embryos harvested from >12 weeks old C57BL/6J pregnant mice (Jackson Laboratory), and subsequently cultured in the same DMEM, no glucose, supplemented with either 25 mM glucose or 10 mM galactose. Cells used for plaque assays – BSC40 and Vero cells – were grown in MEM (Sigma) with 10% FBS, 2 mM L-glutamine, 1X Antibiotic-Antimycotic, and 10 mM HEPES. The following reagents were added at the indicated concentrations unless otherwise noted: 2-deoxyglucose [10 mM (Cayman Chemical Company)], MG132 [1 μM (Sigma)], polybrene [10 μg/mL (Sigma)], universal interferon alpha [1000 U/mL (PBL assay Science)], mouse interferon alpha [1000 U/mL (Invitrogen)], human interferon gamma [1000 U/mL (Invitrogen)], mouse interferon gamma [1000 U/mL (Invitrogen)], feline interferon gamma [1000 U/mL (R&D Systems)]. Viruses used in this study: HSV-1-GFP KOS strain (unpublished) (Dr. David Leib), VACV-Luc-GFP (Dr. Gary Luker)[112], and VSV-Luc (Dr. Sean Whelan)[113]. All phase contrast images were taken with the InvivoGen EVOS^TM^ FL microscope.

### RNA-seq experiment and analysis

A549 cells were grown in glucose or galactose media then pre-treated with either IFN-α or IFN-γ for twenty-four hours followed by mock or vaccinia virus infection for another twenty-four hours. Each infection was performed in triplicate for a total of thirty-six samples. Total RNA was extracted from cells per manufacturer’s instructions with the mirVana miRNA Isolation kit (Ambion, cat. no. AM1561) as it captures small and large RNA species. DNase treatment of the RNA samples was performed using DNA-free DNA Removal Kit (Invitrogen, cat. no. AM1906) per manufacturer’s protocol. RNA library construction was carried out using Ultra II RNA Library Prep Kit for IIIumina (NEB). RNA integrity and library size were assessed with the Agilent tape station (RIN > 6). Concentration was measured by Qubit (A260/A280=1.8-2.2). Sequencing was performed by Genewiz using the Illumina HiSeq 4000 (2x150 bp configuration, single index). The data output was ∼100M raw paired-end reads per sample. Differential analysis was carried out with DESeq2 package by Genewiz [114]. Processed data were filtered by log_2_ fold change ≥ 1 or log_2_ fold change ≤ -1 with adjusted p-value ≤ 0.01 (see supplemental tables). Canonical pathway analysis was performed with QIAGEN Ingenuity Pathway Analysis (QIAGEN IPA) (QIAGEN Inc., https://digitalinsights.qiagen.com/IPA) on genes with log_2_ fold change ≥ 1 or log_2_ fold change ≤ -1 with adjusted p-value ≤ 0.01 from conditions of interest. Data are deposited in NCBI Geo database under accession GSE226242.

### Bioinformatic analysis of data from GTEx portal

Forty-eight defined “core” ISGs were retrieved from published data with a cut-off log_2_ fold change ≥ 1.5 [39]. *HLA* was excluded because it is multi-copy. Thirty-two total human interferons and six interferon receptors (IFNs) were retrieved from The HUGO Gene Nomenclature Committee (HGNC) Gene group reports (https://www.genenames.org/data/genegroup/#!/) for this analysis. Expression levels for the combined set of ISGs, IFNs, and IFN receptors (eighty-six total) genes were analyzed using GTEx Bulk RNA-seq data to determine gene expression levels (median TPM values) across fifty-four tissues. The data were visualized using a cluster heatmap generated with the python seaborn package. *SAT1* was deemed an outlier and excluded from subsequent analysis involving tonic ISG expression. Each of the three gene clusters (high expression, variable expression, and low expression genes) resulting from the cluster analysis was analyzed using core analysis (canonical pathway) in QIAGEN IPA (QIAGEN Inc., https://digitalinsights.qiagen.com/IPA). GTEx tissue specific expression analysis was performed for a subset of genes from each of the three categories for lung, liver, spleen, and whole blood tissues. The GTEx v8 Bulk_RNA-seq and tissue specific RNA-seq data used for the analyses were obtained from the GTEx Portal (https://gtexportal.org/home/datasets) (last accessed on 9/11/2023). Data curation and heatmaps were generated in python 3.8.1 using pandas, NumPy, matplotlib, and seaborn. Pathways bar charts generated using QIAGEN IPA and modified in Adobe Illustrator.

### Plasmids for CRISPR knockout cell lines

*IRF1* and *STAT1* polyclonal KO cell lines were generated using single guide RNAs (sgRNA) cloned into pLentiCRISPR V2 (Addgene #52961) packaged using pSPAX2 and pMD2G. Human *STAT1* and *IRF1* sgRNAs were purchased from GenScript. Two independent sgRNAs were used to generate CRISPR KO cell lines with the lentiviral system.

### Lentiviral generation and transduction

8 x 10^5^ HEK 293T cells were plated per well in a 6-well plate. The next day cells were co-transfected with pSPAX2 (1 μg), pMD2G (1 μg), sgRNA in plentiCRISPR V2 (2 μg) using Lipofectamine 3000 (ThermoFisher, cat. no. L3000008). Supernatant containing lentivirus was collected and pooled together at both 24 and 48 hrs post-transfection from HEK 293T cells. Cellular debris was subsequently removed from pooled supernatants using a 0.45 μm PVDF filter (EMD Millipore). For lentiviral transduction, WT A549 cells were plated in 2x10^5^ cells/well in 6-well plates. The next day, media was removed from A549 cells and transduced with complete DMEM media (1 mL/well) containing viral supernatant (500 μL) and polybrene (10 μg/mL). 18 hours post-transduction media was changed. Puromycin (InvivoGen, 1.5 μg/mL) selection was initiated at 48 hours post-transduction cells and continued for six days before expansion of puromycin-resistant cells. CRISPR KO lines were validated by western blot. gRNA sequences are in S5 Table.

### Western blot analysis

Cells were harvested and lysed in RIPA Lysis and Extraction Buffer (ThermoFisher, cat. no. 89901) supplemented with 1X Halt Protease Inhibitor Cocktail (ThermoFisher, cat. no. 78430), and all samples were incubated on ice for at least 30 minutes. Protein concentration was measured using Bradford assay. Samples were separated by Any kD^TM^ SDS-PAGE (Bio-Rad, cat. no. 4568124) at a constant 110 V for 1 hour. The gel was then wet-transferred to a 0.2 μM Immobilon-PSQ PVDF membrane (Millipore, cat. no. SEQ00010) at 200 mA for 90 min. Membranes were blocked with blocking buffer (5% milk in TBST) for 1 hour at room temperature, and then incubated with primary antibodies at 4°C overnight. The following primary antibodies used in this study were diluted in 1:1000 in 5% milk in TBST: STAT1 rabbit mAB (CST, cat. no. 14994T), phospho-STAT1 rabbit mAB (CST, cat. no. 9167S), IRF1 mAB (CST, cat. no. 8478T), GBP2 pAB (Proteintech, cat no. 11854-1-AP), ISG15 mAB (Santa Cruz Biotech, cat. no. sc-166755), IFITM3 pAB (Proteintech, cat. no. 11714-1-AP), IDO1 pAB (Novus Biologicals, cat. no. NBP1-87703), IFIT1 mAB (CST, cat. no. 14769S), HDAC1 pAB (Proteintech, cat. no. 10197-1-AP), SDHA mAB (CST, cat. no. 11998S), TOM70 pAB (ABclonal, cat. no. A4349), vaccinia virus A27L pAB (BEI resources, cat. no. NR-627), vaccinia virus I3L mAB (a generous gift from Dr. David Evans) [115], HSV-1 VP16 mAB (Santa Cruz Biotech, cat. no. sc-7545), HSV-1 ICP27 mAB (Santa Cruz Biotech, cat. no. sc-69806), PARP mAB (CST, cat. no. 9532S), and β-actin mAB (Sigma, cat. no. A5316). Membranes were washed 3X with TBST and then incubated with secondary antibodies (1:3000 dilution) in 5% milk in TBST for 1 hour at room temperature. Secondary antibodies used for this study were goat anti-rabbit IgG (Bio-Rad, cat. no. 170-6515) and goat anti-mouse IgG (Bio-Rad, cat. no. 170-6516). Membranes were washed three times again with TBST and then developed with Pierce ECL Plus Western Blotting Substrate (ThermoFisher, cat. no. 32132). Blots were imaged using the ChemiDoc MP Imager (Bio-Rad). Densitometry analysis of PARP levels was performed using Image Lab version 6.1 (Bio-Rad). % Cleaved PARP = (cleaved PARP/(Full + Cleaved PARP)) * 100.

### qPCR

Total RNA was extracted from cells using the mirVana miRNA Isolation kit (Ambion, cat. no. AM1561) based on the manufacturer’s protocol. For each sample, 1 μg of total RNA was used for cDNA synthesis using Maxima First-strand cDNA synthesis kit (ThermoFisher, cat. no. R1362). The cDNA synthesis reaction was carried out in a thermocycler (Bio-Rad) – 10 min. at 25°C, 30 min. at 50°C, and termination of enzymatic reaction at 85°C for 5 min. Additional dH_2_O was added to the cDNA for a final 1:5 dilution. 2 μL of diluted cDNA was used as template for each qPCR reaction using Applied Biosystem Power Up SYBR Green Master Mix (ThermoFisher, cat. no. A25776) together with appropriate primers (10 μM/per primer). qPCR was performed in an Applied Biosystems QuantStudio 7 Real-Time PCR instrument following the manufacturer’s instructions. The cycling parameter for qPCR was the following: UDG activation at 50°C for 2 min, followed by activation of dual-lock DNA polymerase at 95°C for 2 min., 40 cycles of 95°C for 15s, 55°C for 19s, 72°C for 1min. Human and mouse *β-actin* were used for controls. Primer sequences are in S5 Table.

### Viral infection

Day 1 (seeding): 2 x 10^5^ A549 cells were seeded per well in 6-well plates in DMEM media, no glucose without sodium pyruvate (Corning, cat.no. 11966025), supplemented with either glucose (25 mM) or galactose (10 mM). Both glucose and galactose supplemented media were made complete with 10% FBS (Gibco, Corning cat. no. 16140071), 2 mM L-glutamine (Corning cat. no. MT25005CI), 10 mM HEPES (Corning, cat.no. 15630080), and 1X Antibiotic-Antimycotic (Gibco, cat.no. 15240112). Day 2 (fresh media added): The next day media was replaced with fresh glucose/galactose supplemented complete media. Day 3 (interferon-priming): Media was replaced with glucose/galactose supplemented complete media with either IFN-α (1000 U/mL) and IFN-γ (1000 U/mL). Day 4 (viral infection): 24 hours post-IFN treatment, cells were infected with viruses at the indicated multiplicity of infections (MOI) – VACV-Luc-GFP (MOI = 0.01 or MOI = 3), VSV-Luc (MOI = 0.01), HSV-1-GFP (MOI = 1) in 1 mL of glucose/galactose supplemented complete media lacking interferon per well for 1 hour at 37°C. After 1 hour incubation, an additional 1 mL of glucose/galactose supplemented complete media lacking interferon was added to each well for a total volume of 2 mL. All assays for a given virus used the same MOI except vaccinia where indicated. Day 5 (assay): Cells were harvested for western blot and qPCR analysis 24 hours post-infection unless otherwise indicated. Viruses for plaque assay analysis were from infections performed on 3 x 10^4^ cells per well in 24-well plates. For Bright-Glo (Promega) luciferase assays, infections were performed on 3 x 10^3^ cells per well in 96-well plates. All assays were performed at either 24 or 48 hours post-infection.

### Plaque assays

The titers of infectious virions for VACV-Luc-GFP and HSV-1-GFP were measured by applying viral inoculums onto BSC40 and Vero cells, respectively. Samples for plaque assays were frozen at 48 hours post-infection and underwent two freeze-thaw cycles at 37°C on the day of assay. 10-fold serial dilutions of thawed samples in MEM were added into 12-well plates (Corning cat. no. 3513) with cell monolayers (2 x 10^5^ cells per well). After 1 hour incubation in 37°C incubator, 1 mL of MEM was added in each well. For HSV-1-GFP plaque assays, an additional 1 mL of overlay containing 1% methylcellulose with MEM was added on each well. The next day, media was aspirated followed by addition of 10% formaldehyde and incubation for 1 hour at room temperature. The cell monolayer was then stained with crystal violet solution (0.1% crystal violet and 20% ethanol) for 1 hour at room temperature to visualize plaques.

### Bright-Glo luciferase assay

3 x 10^3^ A549 cells were plated per well in 100 μL opaque white 96-well plates with clear bottom (Corning, cat. no. 3903). The next day media was replaced with fresh glucose/galactose supplemented complete media. The following day media were removed and cells were treated with 1000 U/mL of IFN-α or IFN-γ in 100 μL of glucose (25 mM) or galactose (10 mM) media. Viral infection was carried out 24 hours post-IFN treatments in media lacking interferons. Bright-Glo luciferase assays were performed at 24 or 48 hours post-infection. 100 μL BrightGlo reagent (Promega, cat no. E2610) was added to each well and incubated at room temperature for 10 minutes. Bioluminescence was measured using BioTek Synergy HTX plate reader.

### IRF1 transient transfection rescue experiment in A549 cells

For the luciferase assay experiment, 3 x 10^3^ *IRF1* knockout cells (sgIRF1_1) were seeded in 100 μL of DMEM glucose (25 mM) media per well on opaque white 96-well plates with a clear bottom (Corning, cat. no. 3903). 200 ng of plasmid was transfected per well using Lipofectamine 3000 (ThermoFisher, cat. no. L3000008) following the manufacturer’s protocol. Three different plasmids were transfected in individual groups: pcDNA3.1, pcDNA3.1 WT single FLAG-human IRF1, and pcDNA3.1 single FLAG-human IRF1 YLP(Y109A/L112A/P113A). All human IRF1 plasmids were CRISPR-resistant through codon optimization and synthesized by Gene Universal. Cells were infected with VACV-Luc-GFP at MOI = 0.01 at 24 hours or 48 hours post-transfection. Bright-Glo luciferase assays were performed, as described above, at 24 hours post-infection. For the western blot experiment, *IRF1* KO cells (sgIRF1_1) were plated 3 x 10^5^ cells per well in 6-well plates in DMEM glucose (25 mM) media. Transfection was carried out with 5 μg of plasmid per well using Lipofectamine 3000. Cells were infected with VACV-Luc-GFP at MOI = 0.01 at 48 hours post-transfection. The lysates were harvested at 24 hours post-infection and subsequently processed for western blot following the method described above.

### Interferon response element (ISRE) luciferase reporter assay

1 x 10^4^ HEK 293T cells were plated per well in 100 μL opaque white 96-well plates with clear bottom (Corning, cat. no 3903). The next day, a total of 200 ng plasmids were co-transfected using Fugene6 Transfection Reagent (Promega, cat. no. E2691). Each group consists of three plasmids. An expression plasmid [pcDNA3.1, pcDNA3.1 WT human IRF1, and pcDNA3.1 human IRF1 YLP (Y109A/L112A/P113A)]; a ISRE reporter [pISRE-Fluc (Agilent Technologies #219092)]; and one pGL4.70 plasmid [hRluc]. Dual-Glo luciferase assays were performed at 48 hours post-transfection. 75μL of Dual-Glo luciferase Assay Reagent (Promega, cat. no. E2920) was added to each well and incubated at room temperature for 10 minutes followed by bioluminescence reading. 75 μL of Dual-Glo Stop & Glo reagent was added to each well and incubated at room temperature for 10 minutes. Bioluminescence was measured using BioTek Synergy HTX plate reader. All human IRF1 plasmids were synthesized by Gene Universal. pGL4.70 [hRluc] was gifted from Dr. Tiffany Reese, UT Southwestern.

### Lactate-Glo assay

A549 cells were plated 3 x 10^3^ per well in 100 μL opaque white 96-well plates with clear bottom (Corning, cat. no. 3903). The next day media was replaced with fresh glucose/galactose supplemented complete media. The following day media were removed, and cells were treated with 1000 U/mL of IFN-α or IFN-γ in 100 μL of glucose (25 mM) or galactose (10 mM) media. 100 μL of media were changed and replenished in each well 24 hours post-IFN treatment. Lactate-Glo assays (Promega, cat no. J5021) were then performed the next day following the manufacturer’s protocol. 50 μL of diluted supernatants (1:1 in PBS) from each well was transferred to a new opaque white 96-well plates, and 50 μL of prepared lactate detection reagent was added to each well and incubated at room temperature for 1 hour. Bioluminescence was measured using BioTek Synergy HTX plate reader.

### ISG time course

Experiments were carried out in A549 cells. Day 1 (seeding): 2 x 10^5^ A549 cells were seeded per well in 6-well plates in DMEM glucose/galactose supplemented complete media. Day 2 (fresh media added): The next day media was replaced with fresh glucose/galactose supplemented complete media. Day 3 (interferon-priming): Media was replaced with glucose/galactose supplemented complete media with IFN-α (1000 U/mL) and IFN-γ (1000 U/mL). RNA and protein were harvested at 4, 8, and 24 hrs post-interferon treatment. For the 48 hr timepoint to match the viral infection protocol, media was replaced 24 hrs post interferon treatment with fresh glucose/galactose supplemented complete media lacking interferons and harvested for RNA/protein analysis 24 hrs thereafter. Membranes were imaged individually to capture differences between glucose and galactose.

### IRF1 target gene analysis

We retrieved known human IRF1 target genes from GSEA Human Gene Sets: GSEA_TFT_IRF1_01 (255 Genes) and GSEA_TFT_IRF1_Q6 (263 Genes), as well as ChEA datasets (502 Genes). These lists were compared against genes upregulated in glucose/IFN-γ and galactose/IFN-γ conditions from RNA-seq. The GSEA datasets were obtained from https://www.gsea-msigdb.org/gsea/msigdb/human/genesets.jsp?collection=TFT and ChEA dataset from (https://maayanlab.cloud/Harmonizome/dataset/CHEA+Transcription+Factor+Targets)[104]. Datasets are in S4 Table.

**S1 Fig. Ingenuity pathway analysis of tonic ISG subclasses.** IPA core canonical pathway analysis was performed on forty-seven core ISGs clustered into three categories (low, high and variable) based on expression patterns across tissues (Fig 1). Dashed line indicates a p-value significance threshold of 0.05.

**S2 Fig. Glucose/galactose with extended interferon-priming does not impact levels of ISG RNAs or non-ISG proteins levels.** (A) Lactate levels for A549 cells grown in glucose/galactose with and without IFN-priming. Assay was performed using Lactate-Glo (N = 4). (B) qPCR of canonical ISGs relative to *β-actin* for cells pre-treated with IFN-γ, grown in glucose or galactose (N = 3). (C) Western blot analysis of non-ISG proteins from different cellular compartments in IFN-primed cells grown in glucose or galactose. Statistical analysis was performed using an unpaired t-test in GraphPad Prism 9.5.1: n.s. not significant, *P ≤ 0.05, ** P < 0.01, *** P ≤ 0.001, **** P ≤ 0.0001. Expression is ordered by the ratio of RNA expression for each gene shown in glucose compared to galactose.

**S3 Fig. Time course of IRF1 and IFITM3 RNA and protein expression across conditions.** A549 cells in glucose or galactose supplemented media were treated with either interferon-α or interferon-γ. RNA and protein samples from the indicated time points (hours: 4,8, 24, 48) were harvested and analyzed for IRF1 and IFITM3 RNA and protein expression. (A) Relative abundance of *IRF1* and *IFITM3* transcripts by qPCR across indicated time points post-IFN treatment in Glc/Gal groups (N=3). (B) Western blot analysis of IRF1 and IFITM3 protein expression at indicated times post-IFN treatment in Glc/Gal. Statistical analysis was performed using an unpaired t-test in GraphPad Prism 9.5.1: n.s. not significant, *P ≤ 0.05, ** P < 0.01, *** P ≤ 0.001, **** P ≤ 0.0001.

**S4 Fig. Characterization of the impact of glucose/galactose on infection outcomes. (**A) Diagrammatic view of experimental set-up in A549 cells and timeline. (B) Vaccinia virus replication (luciferase) assays. Luciferase assays of A549 cells infected with VACV-Luc-GFP grown in either glucose (25 mM) or galactose (10 mM); 24 hours (top) and 48 hours post-infection (bottom). (C) Vaccinia virus replication (luciferase) assays over IFN-γ concentrations. Cells were primed for 24 hours with 1000, 500, 100, 10, or 0 units of IFN-γ followed by infection with VACV-Luc-GFP. (D) Vaccinia virus replication (luciferase) assays over glucose concentrations. Cells were grown in 0, 1, 5, 25 mM of glucose and primed with 1000 units of IFN-γ for 24 hours prior to infection with VACV-Luc-GFP. All cells were infected at MOI = 0.01. (E) HSV-1 viral infections. Qualitative images of plaque assay for cells infected with HSV-1 (MOI = 1) with various dilutions. Glc: glucose (25 mM). Gal: galactose (10 mM). Statistical analysis was performed using an unpaired t-test in GraphPad Prism 9.5.1: n.s. not significant, * P ≤ 0.05, ** P< 0.01, *** P ≤ 0.001, **** P ≤ 0.0001.

**S5 Fig. Vaccinia virus infection (MOI = 3) of A549 cells in media supplemented with glucose/galactose**. A549 cells were infected with vaccinia virus using the same conditions as in Fig 5 and outlined in S4A Fig but with MOI = 3. (A) Vaccinia virus luciferase reporter assays 24 hours post-infection (N = 4). (B) Western blot of vaccinia virus proteins (SSB: vaccinia virus early protein, A27: vaccinia virus late protein). Lysates were harvested at 24 hours post-infection. Glc: glucose (25 mM). Gal: galactose (10 mM). Statistical analysis was performed using an unpaired t-test in GraphPad Prism 9.5.1: n.s. not significant, * P ≤ 0.05, ** P< 0.01, *** P ≤ 0.001, **** P ≤ 0.0001.

**S6 Fig. Analysis of host and interferon-induced responses in glucose/galactose infected with vaccinia virus.** (A) Western blot analysis of total STAT1 and phospho-STAT1 (Tyr 701) in glucose and galactose grown A549 cells, with and without IFN-γ priming, in mock or VACV infected cells. (B) Venn diagram showing the number of differentially expressed genes - log_2_ fold change ≥ 1 or ≤ -1 and adjusted p-value ≤ 0.01 – 24 hrs post mock (orange) and VACV infection (blue) (MOI = 0.01) in cells primed with glucose or galactose-media and pre-treated with IFN-γ prior to infection. Left: genes upregulated in glucose relative to galactose; right: genes upregulated in galactose relative to glucose. (C) IPA core canonical pathway analysis for differentially regulated genes (log_2_ fold change ≥ 1 or ≤ -1 and adjusted p-value ≤ 0.01) shared between mock and VACV infected cells primed with IFN-γ and grown in different carbon sources. Dashed line indicates p-value significant threshold of 0.05. (D) qPCR validation of top gene candidates - from RNA-seq - upregulated in both glucose mock/IFN-γ primed and vaccinia virus/IFN-γ primed infected cells relative to matching conditions but supplemented with galactose; (N = 3). (E) Western blot analysis of top candidate genes. Experiments were carried out according to the timeline outlined in S4A Fig. Statistical analysis was performed using an unpaired t-test in GraphPad Prism 9.5.1: n.s. not significant, * P ≤ 0.05, ** P< 0.01, *** P ≤ 0.001, **** P ≤ 0.0001.

**S7 Fig. Vaccinia virus and VSV infections of *STAT1* and *IRF1* KO cells grown in glucose/galactose with and without IFN-γ priming.** (A) Western blot validation of *STAT1* KO A549 cell lines generated with pLentiCRISPR V2 vectors. Two sgRNAs were used per target gene; sgCtrl – non targeting control. (B) Glucose: VSV-Luc (luciferase) replication assays in *STAT1* KO cells primed with IFN-γ (N = 4). (C) Glucose: VACV-Luc-GFP (luciferase) replication assays in *STAT1* KO cells primed with IFN-γ (N = 4). Experiments were carried out according to the timeline outlined in S4A Fig. Statistical analysis was performed using an unpaired t-test in GraphPad Prism 9.5.1: n.s. not significant, * P ≤ 0.05, ** P< 0.01, *** P ≤ 0.001, **** P ≤ 0.0001.

**S8 Fig. PARP cleavage analysis of sgCtrl and *IRF1* KO cells across conditions with and without vaccinia virus infection.** A549 sgCtrl and *IRF1* KO cells (sgIRF1_1) were infected with vaccinia virus (MOI = 0.01) with or without interferon-priming. Protein lysates were harvested 24 hours post-infection and analyzed by western blot. PARP western blot analysis from (A) mock infected sgCtrl cells, (B) mock infected *IRF1* KO cells, (C) vaccinia infected sgCtrl cells, (D) vaccinia infected *IRF1* KO cells. Experiments were carried out according to the timeline outlined in S4A Fig. VACV: vaccinia virus.

**S9 Fig. Images of sgCtrl and *IRF1* KO cells across conditions with and without vaccinia virus infection.** A549 sgCtrl and *IRF1* KO (sgIRF1_1) cells - cultured in either glucose/galactose media - were infected with vaccinia virus (MOI = 0.01) with or without interferon-priming. Microscope images were taken 24 hours post-infection. (A) Brightfield microscope images at 4X of both mock and vaccinia virus infected sgCtrl cells primed with glucose/galactose and grown with or without IFNs. (B) Brightfield microscope images at 4X of both mock and vaccinia virus infected *IRF1* KO cells primed with glucose/galactose and grown with or without IFNs. (C) Brightfield microscope images at 10X of both mock and vaccinia virus infected sgCtrl cells primed with glucose/galactose and grown with or without IFNs. (D) Brightfield microscope images at 10X of both mock and vaccinia virus infected *IRF1* KO cells primed with glucose/galactose and grown with or without IFNs. Experiments were carried out according to the timeline outlined in S4A Fig. VACV: vaccinia virus.

**S10 Fig. qPCR of vaccinia virus RNAs *IRF1* KO cells: glucose with and without IFN-γ priming.** qPCR of VACV transcripts – early (*I3L, J2R*) and late (*F17R, K3L*) genes in sgCtrl, *IRF1* KO (sgIRF1_1) cells, and sgIRF1_2 where *IRF1* KO what is inefficient in glucose/IFN-γ. (N = 3). Statistical analysis was performed using an unpaired t-test in GraphPad Prism 9.5.1: n.s. not significant, * P ≤ 0.05, ** P< 0.01, *** P ≤ 0.001, **** P ≤ 0.0001.

**S11 Fig. IRF1 Target Genes Analysis.** To explore whether there was an IRF1 target gene signature for glucose/IFN-γ cells relative to galactose/IFN-γ cells, we examined differentially expressed genes between the two conditions for evidence of IRF1 regulation. Analysis of three available IRF1 target gene sets, which consist of predicted targets and targets identified from -omics studies (see methods), identified only a limited number of “IRF1 target genes” that differed between our RNA-seq of glucose/IFN-γ cells and galactose/IFN-γ cells. Integrated IRF1 target gene analysis of RNA-seq dataset of glucose/IFN-γ and galactose/IFN-γ cells is shown. Genes highlighted in purple are present under both mock and VACV infected conditions. Genes highlighted in orange are present under only mock-infected conditions. Glucose-upregulated genes in IFN-γ are colored in blue. Galactose-upregulated genes in IFN-γ are colored in blue. See S4 Table.

**S1 Table Related to Fig 1**, **Fig 2 and S1 Fig**

Sheet related to Fig 1:

Median log_2_TPM (transcripts per million) RNA-Seq values for forty-eight human core ISGs and thirty-eight human interferons/interferon receptors across fifty-four tissues. *HLA* was excluded as it is a multi-copy gene. RNA-Seq data were retrieved from GTEx Analysis V8 (https://gtexportal.org/home/datasets). Core ISG designation from [39]. Interferons and interferon receptors were retrieved from The HUGO Gene Nomenclature Committee (HGNC) Gene group reports (https://www.genenames.org/data/genegroup/#!/).

Sheets related to Fig 2:

Log_2_TPM (transcripts per million) RNA-Seq values and statistics for representatives from ISG subclasses - 2 high, 2 low, and 5 variable - across 578 human lung tissue samples, 226 human liver tissue samples, 241 human spleen tissue samples, and 755 human whole blood tissue samples. RNA-Seq data were retrieved from GTEx Analysis V8 (https://gtexportal.org/home/datasets). Core ISG designation is from [39]. Statistics table was generated using python pandas library (https://pandas.pydata.org/).

Sheet related to S1 Fig:

Gene list used for pathway analysis of tonic ISG subclasses. Total of forty-seven core ISG genes from Fig 1. Average RNA-Seq log_2_ fold-change were obtained from [39]. *SAT1* was deemed an outlier and excluded along with interferons and interferon receptors. Core ISGs are classified into three clusters based on their expression level across tissues. Pathway analysis was performed using QIAGEN IPA (QIAGEN Inc., https://digitalinsights.qiagen.com/IPA) with ID and log_2_ fold-change expression values. ISG – interferon-stimulated gene

**S2 Table related to S6 Fig**

Sheet related to S6B Fig GlucoseUpregulatedGenes:

List of cellular genes (with RNA-Seq values) upregulated in glucose media and IFN-γ treated relative to galactose and IFN-γ treated. List generated by comparison of differentially expressed genes (DEGs) from 1) glucose/galactose A549 cultured cells treated with IFN-γ and mock infected with 2) glucose/galactose A549 cultured cells treated with IFN-γ and infected with vaccinia virus (MOI = 0.01). RNA-Seq data were curated with the cut-off values of log_2_ fold-change ≥ 1 or log_2_ fold-change ≤ -1 with an adjusted p-value ≤ 0.01 under mock infection condition. Given the direction of comparison, positive log_2_ fold-change values indicate cellular genes upregulated in galactose, and negative log_2_ fold-change values indicate cellular genes upregulated in glucose.

Sheet related to S6B Fig GalactoseUpregulatedGenes:

List of cellular genes (with RNA-Seq values) upregulated in galactose media and IFN-γ treated relative to glucose media and IFN-γ treated. List obtained from comparison of DEGs from 1) glucose/galactose A549 cultured cells treated with IFN-γ and mock infected with 2) glucose/galactose A549 cultured cells treated with IFN-γ and infected with vaccinia virus (MOI = 0.01). RNA-Seq data were curated with the cut-off values of log_2_ fold-change ≥ 1 or log_2_ fold-change ≤ -1 with an adjusted p-value ≤ 0.01 under mock infection condition. Given the direction of comparison, positive log_2_ fold-change values indicate cellular genes upregulated in galactose, and negative log_2_ fold-change values indicate cellular genes upregulated in glucose.

**S3 Table related to S6C Fig**

Sheet related to S6C Fig GlucoseUpregulated:

Gene list (with RNA-Seq values) used for pathway analysis in S6C Fig. Cellular genes upregulated in both glucose/IFN-γ/mock infected and glucose/IFN-γ/vaccinia virus-infected relative to matching conditions but in galactose media. Comparison made using DEGs from 1) glucose/galactose A549 cultured cells treated with IFN-γ and mock infected with 2) glucose/galactose A549 cultured cells treated with IFN-γ and infected with vaccinia virus (MOI = 0.01). RNA-Seq data were curated with the cut-off values of log_2_ fold-change ≥ 1 or log_2_ fold-change ≤ -1 with an adjusted p-value ≤ 0.01 under mock infection condition. Data were analyzed with QIAGEN Ingenuity Pathway Analysis (QIAGEN IPA).

Sheet related to S6C Fig GalactoseUpregulated:

Gene list (with RNA-Seq values) used for pathway analysis in S6C Fig. Cellular genes upregulated in both galactose/IFN-γ/mock infected and galactose/IFN-γ/vaccinia virus infected relative to matching conditions but in glucose media. Comparison made using DEGs from 1) glucose/galactose A549 cultured cells treated with IFN-γ and mock infected with 2) glucose/galactose A549 cultured cells treated with IFN-γ and infected with vaccinia virus (M.O.I. = 0.01). RNA-Seq data were curated with the cut-off values of log_2_ fold-change ≥ 1 or log_2_ fold-change ≤ -1 with an adjusted p-value ≤ 0.01 under mock infection condition. Data were analyzed with QIAGEN Ingenuity Pathway Analysis (QIAGEN IPA).

**S4 Table related to S11 Fig**

Gene lists used for IRF1 target gene analysis in S11 Fig.

IRF1 target gene lists obtained from GSEA datasets:

https://www.gsea-msigdb.org/gsea/msigdb/human/geneset/IRF1_01.html https://www.gsea-msigdb.org/gsea/msigdb/human/geneset/IRF1_Q6.html

ChEA Dataset https://maayanlab.cloud/Harmonizome/gene_set/IRF1/CHEA+Transcription+Factor+Targets

Comparisons were made using 1) list of cellular genes (with RNA-Seq values) upregulated in glucose media and IFN-γ treated conditions and 2) list of cellular genes (with RNA-Seq values) upregulated in galactose media and IFN-γ treated conditions. Given the direction of comparison, positive log_2_ fold-change values indicate cellular genes upregulated in galactose, and negative log_2_ fold-change values indicate cellular genes upregulated in glucose.

**S5 Table**

Sequences for primers and guide RNAs used in this study.

## ACKNOWLEDGEMENTS

We thank Dr. Steve Baker, and John McCormick for comments on the manuscript as well as Dr. Tiffany Reese for many hallway conversations. We also express our gratitude to Dr. Don Gammon, Dr. Qing Zhang, Dr. Sean Whelan, Dr. Gary Luker, and Dr. David Leib for sharing reagents with us for this work. This study in the Hancks lab was supported in part, by R00 GM119126-04, 1R35GM142689-01, and a Recruitment of First-Time, Tenure-Track Faculty Award from the Cancer Prevention & Research Institute of Texas (RR 170047) to D.C.H. T.C. was supported, in part, by a National Institutes of Health by Immunology Training Grant No. 2T32AI005284-41A1.

## AUTHOR CONTRIBUTIONS

Conceptualization, T.C. and D.C.H.; Data curation, T.C., J.A., S.C., D.C.H.; Validation, T.C., J.A., S.C., D.C.H.; Investigation, T.C., J.A., S.C., D.C.H.; Methodology, T.C., J.A., S.C., S.C., M.S., R.C.O., D.C.H.; Formal analysis, T.C., J.A., S.C., D.C.H.; Project administration, D.C.H.; Writing – Original Draft, T.C. and D.C.H.; Writing – Review & Editing, T.C., J.A., S.C., M.S., D.C.H.; Visualization, T.C., J.A., S.C., and D.C.H.; Supervision, D.C.H.; Funding Acquisition, D.C.H.

## REFERENCES

1. Liston A, Humblet-Baron S, Duffy D, Goris A. Human immune diversity: from evolution to modernity. Nat Immunol. 2021;22: 1479–1489. doi:10.1038/s41590-021-01058-1

2. Dean M, Carrington M, Winkler C, Huttley GA, Smith MW, Allikmets R, et al. Genetic restriction of HIV-1 infection and progression to AIDS by a deletion allele of the CKR5 structural gene. Hemophilia Growth and Development Study, Multicenter AIDS Cohort Study, Multicenter Hemophilia Cohort Study, San Francisco City Cohort, ALIVE Study. Science. 1996;273: 1856–1862. doi:10.1126/science.273.5283.1856

3. Liu R, Paxton WA, Choe S, Ceradini D, Martin SR, Horuk R, et al. Homozygous defect in HIV-1 coreceptor accounts for resistance of some multiply-exposed individuals to HIV-1 infection. Cell. 1996;86: 367–377. doi:10.1016/s0092-8674(00)80110-5

4. Samson M, Libert F, Doranz BJ, Rucker J, Liesnard C, Farber CM, et al. Resistance to HIV-1 infection in caucasian individuals bearing mutant alleles of the CCR-5 chemokine receptor gene. Nature. 1996;382: 722–725. doi:10.1038/382722a0

5. Sironi M, Cagliani R, Forni D, Clerici M. Evolutionary insights into host-pathogen interactions from mammalian sequence data. Nat Rev Genet. 2015;16: 224–236. doi:10.1038/nrg3905

6. Daugherty MD, Malik HS. Rules of engagement: molecular insights from host-virus arms races. Annu Rev Genet. 2012;46: 677–700. doi:10.1146/annurev-genet-110711-155522

7. Akalu YT, Bogunovic D. Inborn errors of immunity: an expanding universe of disease and genetic architecture. Nat Rev Genet. 2024;25: 184–195. doi:10.1038/s41576-023-00656-z

8. Russell CD, Lone NI, Baillie JK. Comorbidities, multimorbidity and COVID-19. Nat Med. 2023;29: 334–343. doi:10.1038/s41591-022-02156-9

9. Rouse BT, Sehrawat S. Immunity and immunopathology to viruses: what decides the outcome? Nat Rev Immunol. 2010;10: 514–526. doi:10.1038/nri2802

10. Hansen JD, Vojtech LN, Laing KJ. Sensing disease and danger: a survey of vertebrate PRRs and their origins. Dev Comp Immunol. 2011;35: 886–897. doi:10.1016/j.dci.2011.01.008

11. Secombes CJ, Zou J. Evolution of Interferons and Interferon Receptors. Front Immunol. 2017;8: 209. doi:10.3389/fimmu.2017.00209

12. Sun L, Wu J, Du F, Chen X, Chen ZJ. Cyclic GMP-AMP synthase is a cytosolic DNA sensor that activates the type I interferon pathway. Science. 2013;339: 786–791. doi:10.1126/science.1232458

13. Ablasser A, Hur S. Regulation of cGAS- and RLR-mediated immunity to nucleic acids. Nat Immunol. 2020;21: 17–29. doi:10.1038/s41590-019-0556-1

14. de Oliveira Mann CC, Hornung V. Molecular mechanisms of nonself nucleic acid recognition by the innate immune system. Eur J Immunol. 2021;51: 1897–1910. doi:10.1002/eji.202049116

15. Fitzgerald KA, Kagan JC. Toll-like Receptors and the Control of Immunity. Cell. 2020;180: 1044–1066. doi:10.1016/j.cell.2020.02.041

16. Wu J, Sun L, Chen X, Du F, Shi H, Chen C, et al. Cyclic GMP-AMP is an endogenous second messenger in innate immune signaling by cytosolic DNA. Science. 2013;339: 826–830. doi:10.1126/science.1229963

17. Schoggins JW. Interferon-Stimulated Genes: What Do They All Do? Annu Rev Virol. 2019;6: 567–584. doi:10.1146/annurev-virology-092818-015756

18. Platanias LC. Mechanisms of type-I- and type-II-interferon-mediated signalling. Nat Rev Immunol. 2005;5: 375–386. doi:10.1038/nri1604

19. Der SD, Zhou A, Williams BR, Silverman RH. Identification of genes differentially regulated by interferon alpha, beta, or gamma using oligonucleotide arrays. Proc Natl Acad Sci U S A. 1998;95: 15623–15628. doi:10.1073/pnas.95.26.15623

20. Stetson DB, Medzhitov R. Type I interferons in host defense. Immunity. 2006;25: 373–381. doi:10.1016/j.immuni.2006.08.007

21. Ivashkiv LB. IFNγ: signalling, epigenetics and roles in immunity, metabolism, disease and cancer immunotherapy. Nat Rev Immunol. 2018;18: 545–558. doi:10.1038/s41577-018-0029-z

22. Lazear HM, Schoggins JW, Diamond MS. Shared and Distinct Functions of Type I and Type III Interferons. Immunity. 2019;50: 907–923. doi:10.1016/j.immuni.2019.03.025

23. Philips RL, Wang Y, Cheon H, Kanno Y, Gadina M, Sartorelli V, et al. The JAK-STAT pathway at 30: Much learned, much more to do. Cell. 2022;185: 3857–3876. doi:10.1016/j.cell.2022.09.023

24. Darnell JEJ, Kerr IM, Stark GR. Jak-STAT pathways and transcriptional activation in response to IFNs and other extracellular signaling proteins. Science. 1994;264: 1415– 1421. doi:10.1126/science.8197455

25. Crow YJ, Stetson DB. The type I interferonopathies: 10 years on. Nat Rev Immunol. 2022;22: 471–483. doi:10.1038/s41577-021-00633-9

26. Crow YJ, Hayward BE, Parmar R, Robins P, Leitch A, Ali M, et al. Mutations in the gene encoding the 3’-5’ DNA exonuclease TREX1 cause Aicardi-Goutières syndrome at the AGS1 locus. Nat Genet. 2006;38: 917–920. doi:10.1038/ng1845

27. Ferreira RC, Guo H, Coulson RMR, Smyth DJ, Pekalski ML, Burren OS, et al. A type I interferon transcriptional signature precedes autoimmunity in children genetically at risk for type 1 diabetes. Diabetes. 2014;63: 2538–2550. doi:10.2337/db13-1777

28. Chiche L, Jourde-Chiche N, Whalen E, Presnell S, Gersuk V, Dang K, et al. Modular transcriptional repertoire analyses of adults with systemic lupus erythematosus reveal distinct type I and type II interferon signatures. Arthritis Rheumatol Hoboken NJ. 2014;66: 1583–1595. doi:10.1002/art.38628

29. Lee D, Le Pen J, Yatim A, Dong B, Aquino Y, Ogishi M, et al. Inborn errors of OAS-RNase L in SARS-CoV-2-related multisystem inflammatory syndrome in children. Science. 2022; eabo3627. doi:10.1126/science.abo3627

30. Warren EB, Gordon-Lipkin EM, Cheung F, Chen J, Mukherjee A, Apps R, et al. Inflammatory and interferon gene expression signatures in patients with mitochondrial disease. J Transl Med. 2023;21: 331. doi:10.1186/s12967-023-04180-w

31. Zhang W, Wang G, Xu Z-G, Tu H, Hu F, Dai J, et al. Lactate Is a Natural Suppressor of RLR Signaling by Targeting MAVS. Cell. 2019;178: 176–189.e15. doi:10.1016/j.cell.2019.05.003

32. Sheehy AM, Gaddis NC, Malim MH. The antiretroviral enzyme APOBEC3G is degraded by the proteasome in response to HIV-1 Vif. Nat Med. 2003;9: 1404–1407. doi:10.1038/nm945

33. García-Sastre A. Ten Strategies of Interferon Evasion by Viruses. Cell Host Microbe. 2017;22: 176–184. doi:10.1016/j.chom.2017.07.012

34. Mostafavi S, Yoshida H, Moodley D, LeBoité H, Rothamel K, Raj T, et al. Parsing the Interferon Transcriptional Network and Its Disease Associations. Cell. 2016;164: 564–578. doi:10.1016/j.cell.2015.12.032

35. Wu X, Dao Thi VL, Huang Y, Billerbeck E, Saha D, Hoffmann H-H, et al. Intrinsic Immunity Shapes Viral Resistance of Stem Cells. Cell. 2018;172: 423–438.e25. doi:10.1016/j.cell.2017.11.018

36. Abt MC, Osborne LC, Monticelli LA, Doering TA, Alenghat T, Sonnenberg GF, et al. Commensal bacteria calibrate the activation threshold of innate antiviral immunity. Immunity. 2012;37: 158–170. doi:10.1016/j.immuni.2012.04.011

37. The GTEx Consortium. The human transcriptome across tissues and individuals. Science. 2015;348: 660–665. doi:10.1126/science.aaa0355

38. Human genomics. The Genotype-Tissue Expression (GTEx) pilot analysis: multitissue gene regulation in humans. Science. 2015;348: 648–660. doi:10.1126/science.1262110

39. Shaw AE, Hughes J, Gu Q, Behdenna A, Singer JB, Dennis T, et al. Fundamental properties of the mammalian innate immune system revealed by multispecies comparison of type I interferon responses. PLoS Biol. 2017;15: e2004086. doi:10.1371/journal.pbio.2004086

40. Carter B, Zhao K. The epigenetic basis of cellular heterogeneity. Nat Rev Genet. 2021;22: 235–250. doi:10.1038/s41576-020-00300-0

41. Armingol E, Officer A, Harismendy O, Lewis NE. Deciphering cell-cell interactions and communication from gene expression. Nat Rev Genet. 2021;22: 71–88. doi:10.1038/s41576-020-00292-x

42. Dai Z, Ramesh V, Locasale JW. The evolving metabolic landscape of chromatin biology and epigenetics. Nat Rev Genet. 2020;21: 737–753. doi:10.1038/s41576-020-0270-8

43. Palmer CS. Innate metabolic responses against viral infections. Nat Metab. 2022;4: 1245– 1259. doi:10.1038/s42255-022-00652-3

44. Pant A, Dsouza L, Yang Z. Alteration in Cellular Signaling and Metabolic Reprogramming during Viral Infection. mBio. 2021;12: e0063521. doi:10.1128/mBio.00635-21

45. Polcicova K, Badurova L, Tomaskova J. Metabolic reprogramming as a feast for virus replication. Acta Virol. 2020;64: 201–215. doi:10.4149/av_2020_210

46. Sanchez EL, Lagunoff M. Viral activation of cellular metabolism. Virology. 2015;479–480: 609–618. doi:10.1016/j.virol.2015.02.038

47. Kominsky DJ, Campbell EL, Colgan SP. Metabolic shifts in immunity and inflammation. J Immunol. 2010;184: 4062–4068. doi:10.4049/jimmunol.0903002

48. Ghazarian M, Revelo XS, Nøhr MK, Luck H, Zeng K, Lei H, et al. Type I Interferon Responses Drive Intrahepatic T cells to Promote Metabolic Syndrome. Sci Immunol. 2017;2: eaai7616. doi:10.1126/sciimmunol.aai7616

49. Bornstein SR, Dalan R, Hopkins D, Mingrone G, Boehm BO. Endocrine and metabolic link to coronavirus infection. Nat Rev Endocrinol. 2020;16: 297–298. doi:10.1038/s41574-020-0353-9

50. Stefan N. Metabolic disorders, COVID-19 and vaccine-breakthrough infections. Nat Rev Endocrinol. 2022;18: 75–76. doi:10.1038/s41574-021-00608-9

51. Liu X, Kim CN, Yang J, Jemmerson R, Wang X. Induction of apoptotic program in cell-free extracts: requirement for dATP and cytochrome c. Cell. 1996;86: 147–157. doi:10.1016/s0092-8674(00)80085-9

52. West AP, Brodsky IE, Rahner C, Woo DK, Erdjument-Bromage H, Tempst P, et al. TLR signalling augments macrophage bactericidal activity through mitochondrial ROS. Nature. 2011;472: 476–480. doi:10.1038/nature09973

53. Weindel CG, Martinez EL, Zhao X, Mabry CJ, Bell SL, Vail KJ, et al. Mitochondrial ROS promotes susceptibility to infection via gasdermin D-mediated necroptosis. Cell. 2022;185: 3214–3231.e23. doi:10.1016/j.cell.2022.06.038

54. Sorouri M, Chang T, Jesudhasan P, Pinkham C, Elde NC, Hancks DC. Signatures of host-pathogen evolutionary conflict reveal MISTR-A conserved MItochondrial STress Response network. PLoS Biol. 2020;18: e3001045. doi:10.1371/journal.pbio.3001045

55. Sorouri M, Chang T, Hancks DC. Mitochondria and Viral Infection: Advances and Emerging Battlefronts. mBio. 2022;13: e0209621. doi:10.1128/mbio.02096-21

56. Weinberg F, Hamanaka R, Wheaton WW, Weinberg S, Joseph J, Lopez M, et al. Mitochondrial metabolism and ROS generation are essential for Kras-mediated tumorigenicity. Proc Natl Acad Sci U S A. 2010;107: 8788–8793. doi:10.1073/pnas.1003428107

57. Marroquin LD, Hynes J, Dykens JA, Jamieson JD, Will Y. Circumventing the Crabtree effect: replacing media glucose with galactose increases susceptibility of HepG2 cells to mitochondrial toxicants. Toxicol Sci Off J Soc Toxicol. 2007;97: 539–547. doi:10.1093/toxsci/kfm052

58. O’Neill LAJ, Kishton RJ, Rathmell J. A guide to immunometabolism for immunologists. Nat Rev Immunol. 2016;16: 553–565. doi:10.1038/nri.2016.70

59. Rodríguez-Prados J-C, Través PG, Cuenca J, Rico D, Aragonés J, Martín-Sanz P, et al. Substrate fate in activated macrophages: a comparison between innate, classic, and alternative activation. J Immunol Baltim Md 1950. 2010;185: 605–614. doi:10.4049/jimmunol.0901698

60. Hernaez B, Alcamí A. Virus-encoded cytokine and chemokine decoy receptors. Curr Opin Immunol. 2020;66: 50–56. doi:10.1016/j.coi.2020.04.008

61. Colamonici OR, Domanski P, Sweitzer SM, Larner A, Buller RM. Vaccinia virus B18R gene encodes a type I interferon-binding protein that blocks interferon alpha transmembrane signaling. J Biol Chem. 1995;270: 15974–15978. doi:10.1074/jbc.270.27.15974

62. Li Y, Banerjee S, Goldstein SA, Dong B, Gaughan C, Rath S, et al. Ribonuclease L mediates the cell-lethal phenotype of double-stranded RNA editing enzyme ADAR1 deficiency in a human cell line. eLife. 2017;6: e25687. doi:10.7554/eLife.25687

63. Schoggins JW, Wilson SJ, Panis M, Murphy MY, Jones CT, Bieniasz P, et al. A diverse range of gene products are effectors of the type I interferon antiviral response. Nature. 2011;472: 481–485. doi:10.1038/nature09907

64. Lu M, Yang C, Li M, Yi Q, Lu G, Wu Y, et al. A conserved interferon regulation factor 1 (IRF-1) from Pacific oyster Crassostrea gigas functioned as an activator of IFN pathway. Fish Shellfish Immunol. 2018;76: 68–77. doi:10.1016/j.fsi.2018.02.024

65. Yu H, Bruneau RC, Brennan G, Rothenburg S. Battle Royale: Innate Recognition of Poxviruses and Viral Immune Evasion. Biomedicines. 2021;9. doi:10.3390/biomedicines9070765

66. Bratke KA, McLysaght A, Rothenburg S. A survey of host range genes in poxvirus genomes. Infect Genet Evol J Mol Epidemiol Evol Genet Infect Dis. 2013;14: 406–425. doi:10.1016/j.meegid.2012.12.002

67. Terajima M, Leporati AM. Role of Indoleamine 2,3-Dioxygenase in Antiviral Activity of Interferon-gamma Against Vaccinia Virus. Viral Immunol. 2005;18: 722–729. doi:10.1089/vim.2005.18.722

68. Trilling M, Le VTK, Zimmermann A, Ludwig H, Pfeffer K, Sutter G, et al. Gamma interferon-induced interferon regulatory factor 1-dependent antiviral response inhibits vaccinia virus replication in mouse but not human fibroblasts. J Virol. 2009;83: 3684– 3695. doi:10.1128/JVI.02042-08

69. Meng X, Schoggins J, Rose L, Cao J, Ploss A, Rice CM, et al. C7L family of poxvirus host range genes inhibits antiviral activities induced by type I interferons and interferon regulatory factor 1. J Virol. 2012;86: 4538–4547. doi:10.1128/JVI.06140-11

70. Hutchens M, Luker KE, Sottile P, Sonstein J, Lukacs NW, Núñez G, et al. TLR3 increases disease morbidity and mortality from vaccinia infection. J Immunol Baltim Md 1950. 2008;180: 483–491. doi:10.4049/jimmunol.180.1.483

71. Moss B. Poxvirus DNA replication. Cold Spring Harb Perspect Biol. 2013;5. doi:10.1101/cshperspect.a010199

72. Kimura T, Nakayama K, Penninger J, Kitagawa M, Harada H, Matsuyama T, et al. Involvement of the IRF-1 transcription factor in antiviral responses to interferons. Science. 1994;264: 1921–1924. doi:10.1126/science.8009222

73. Forero A, Ozarkar S, Li H, Lee CH, Hemann EA, Nadjsombati MS, et al. Differential Activation of the Transcription Factor IRF1 Underlies the Distinct Immune Responses Elicited by Type I and Type III Interferons. Immunity. 2019;51: 451–464.e6. doi:10.1016/j.immuni.2019.07.007

74. Frontini M, Vijayakumar M, Garvin A, Clarke N. A ChIP-chip approach reveals a novel role for transcription factor IRF1 in the DNA damage response. Nucleic Acids Res. 2009;37: 1073–1085. doi:10.1093/nar/gkn1051

75. Dornan D, Eckert M, Wallace M, Shimizu H, Ramsay E, Hupp TR, et al. Interferon regulatory factor 1 binding to p300 stimulates DNA-dependent acetylation of p53. Mol Cell Biol. 2004;24: 10083–10098. doi:10.1128/MCB.24.22.10083-10098.2004

76. Liu S-Y, Sanchez DJ, Aliyari R, Lu S, Cheng G. Systematic identification of type I and type II interferon-induced antiviral factors. Proc Natl Acad Sci U S A. 2012;109: 4239–4244. doi:10.1073/pnas.1114981109

77. Wickenhagen A, Sugrue E, Lytras S, Kuchi S, Noerenberg M, Turnbull ML, et al. A prenylated dsRNA sensor protects against severe COVID-19. Science. 2021;374: eabj3624. doi:10.1126/science.abj3624

78. de Veer MJ, Holko M, Frevel M, Walker E, Der S, Paranjape JM, et al. Functional classification of interferon-stimulated genes identified using microarrays. J Leukoc Biol. 2001;69: 912–920.

79. Samarajiwa SA, Forster S, Auchettl K, Hertzog PJ. INTERFEROME: the database of interferon regulated genes. Nucleic Acids Res. 2009;37: D852–857. doi:10.1093/nar/gkn732

80. Marié IJ, Brambilla L, Azzouz D, Chen Z, Baracho GV, Arnett A, et al. Tonic interferon restricts pathogenic IL-17-driven inflammatory disease via balancing the microbiome. eLife. 2021;10. doi:10.7554/eLife.68371

81. Veyer DL, Carrara G, Maluquer de Motes C, Smith GL. Vaccinia virus evasion of regulated cell death. Immunol Lett. 2017;186: 68–80. doi:10.1016/j.imlet.2017.03.015

82. Wang G, Zhang D, Orchard RC, Hancks DC, Reese TA. Norovirus MLKL-like protein initiates cell death to induce viral egress. Nature. 2023;616: 152–158. doi:10.1038/s41586-023-05851-w

83. Brass AL, Huang I-C, Benita Y, John SP, Krishnan MN, Feeley EM, et al. The IFITM proteins mediate cellular resistance to influenza A H1N1 virus, West Nile virus, and dengue virus. Cell. 2009;139: 1243–1254. doi:10.1016/j.cell.2009.12.017

84. Everitt AR, Clare S, Pertel T, John SP, Wash RS, Smith SE, et al. IFITM3 restricts the morbidity and mortality associated with influenza. Nature. 2012;484: 519–523. doi:10.1038/nature10921

85. Shi G, Kenney AD, Kudryashova E, Zani A, Zhang L, Lai KK, et al. Opposing activities of IFITM proteins in SARS-CoV-2 infection. EMBO J. 2021;40: e106501. doi:10.15252/embj.2020106501

86. Scherer PE, Kirwan JP, Rosen CJ. Post-acute sequelae of COVID-19: A metabolic perspective. eLife. 2022;11. doi:10.7554/eLife.78200

87. Smith M, Honce R, Schultz-Cherry S. Metabolic Syndrome and Viral Pathogenesis: Lessons from Influenza and Coronaviruses. J Virol. 2020;94. doi:10.1128/JVI.00665-20

88. Perakakis N, Harb H, Hale BG, Varga Z, Steenblock C, Kanczkowski W, et al. Mechanisms and clinical relevance of the bidirectional relationship of viral infections with metabolic diseases. Lancet Diabetes Endocrinol. 2023;11: 675–693. doi:10.1016/S2213-8587(23)00154-7

89. Liu Z, Nailwal H, Rector J, Rahman MM, Sam R, McFadden G, et al. A class of viral inducer of degradation of the necroptosis adaptor RIPK3 regulates virus-induced inflammation. Immunity. 2021;54: 247–258.e7. doi:10.1016/j.immuni.2020.11.020

90. Herbert MH, Squire CJ, Mercer AA. Poxviral ankyrin proteins. Viruses. 2015;7: 709–738. doi:10.3390/v7020709

91. Goffinet C, Allespach I, Homann S, Tervo H-M, Habermann A, Rupp D, et al. HIV-1 antagonism of CD317 is species specific and involves Vpu-mediated proteasomal degradation of the restriction factor. Cell Host Microbe. 2009;5: 285–297. doi:10.1016/j.chom.2009.01.009

92. Arroyo JD, Jourdain AA, Calvo SE, Ballarano CA, Doench JG, Root DE, et al. A Genome-wide CRISPR Death Screen Identifies Genes Essential for Oxidative Phosphorylation. Cell Metab. 2016;24: 875–885. doi:10.1016/j.cmet.2016.08.017

93. van der Windt GJW, Everts B, Chang C-H, Curtis JD, Freitas TC, Amiel E, et al. Mitochondrial respiratory capacity is a critical regulator of CD8+ T cell memory development. Immunity. 2012;36: 68–78. doi:10.1016/j.immuni.2011.12.007

94. Kawai T, Takahashi K, Sato S, Coban C, Kumar H, Kato H, et al. IPS-1, an adaptor triggering RIG-I- and Mda5-mediated type I interferon induction. Nat Immunol. 2005;6: 981–988. doi:10.1038/ni1243

95. Seth RB, Sun L, Ea C-K, Chen ZJ. Identification and characterization of MAVS, a mitochondrial antiviral signaling protein that activates NF-kappaB and IRF 3. Cell. 2005;122: 669–682. doi:10.1016/j.cell.2005.08.012

96. Mills EL, Kelly B, O’Neill LAJ. Mitochondria are the powerhouses of immunity. Nat Immunol. 2017;18: 488–498. doi:10.1038/ni.3704

97. Tannahill GM, Curtis AM, Adamik J, Palsson-McDermott EM, McGettrick AF, Goel G, et al. Succinate is an inflammatory signal that induces IL-1β through HIF-1α. Nature. 2013;496: 238–242. doi:10.1038/nature11986

98. West AP, Shadel GS, Ghosh S. Mitochondria in innate immune responses. Nat Rev Immunol. 2011;11: 389–402. doi:10.1038/nri2975

99. Yoneyama M, Kikuchi M, Natsukawa T, Shinobu N, Imaizumi T, Miyagishi M, et al. The RNA helicase RIG-I has an essential function in double-stranded RNA-induced innate antiviral responses. Nat Immunol. 2004;5: 730–737. doi:10.1038/ni1087

100. Meylan E, Curran J, Hofmann K, Moradpour D, Binder M, Bartenschlager R, et al. Cardif is an adaptor protein in the RIG-I antiviral pathway and is targeted by hepatitis C virus. Nature. 2005;437: 1167–1172. doi:10.1038/nature04193

101. Mills EL, Ryan DG, Prag HA, Dikovskaya D, Menon D, Zaslona Z, et al. Itaconate is an anti-inflammatory metabolite that activates Nrf2 via alkylation of KEAP1. Nature. 2018;556: 113–117. doi:10.1038/nature25986

102. Wang A, Huen SC, Luan HH, Yu S, Zhang C, Gallezot J-D, et al. Opposing Effects of Fasting Metabolism on Tissue Tolerance in Bacterial and Viral Inflammation. Cell. 2016;166: 1512–1525.e12. doi:10.1016/j.cell.2016.07.026

103. Price JV, Russo D, Ji DX, Chavez RA, DiPeso L, Lee AY-F, et al. IRG1 and Inducible Nitric Oxide Synthase Act Redundantly with Other Interferon-Gamma-Induced Factors To Restrict Intracellular Replication of Legionella pneumophila. mBio. 2019;10. doi:10.1128/mBio.02629-19

104. Dai A, Cao S, Dhungel P, Luan Y, Liu Y, Xie Z, et al. Ribosome Profiling Reveals Translational Upregulation of Cellular Oxidative Phosphorylation mRNAs during Vaccinia Virus-Induced Host Shutoff. J Virol. 2017;91. doi:10.1128/JVI.01858-16

105. Pant A, Dsouza L, Cao S, Peng C, Yang Z. Viral growth factor- and STAT3 signaling-dependent elevation of the TCA cycle intermediate levels during vaccinia virus infection. PLoS Pathog. 2021;17: e1009303. doi:10.1371/journal.ppat.1009303

106. Greseth MD, Traktman P. De novo fatty acid biosynthesis contributes significantly to establishment of a bioenergetically favorable environment for vaccinia virus infection. PLoS Pathog. 2014;10: e1004021. doi:10.1371/journal.ppat.1004021

107. Pant A, Cao S, Yang Z. Asparagine Is a Critical Limiting Metabolite for Vaccinia Virus Protein Synthesis during Glutamine Deprivation. J Virol. 2019;93. doi:10.1128/JVI.01834-18

108. Fontaine KA, Camarda R, Lagunoff M. Vaccinia virus requires glutamine but not glucose for efficient replication. J Virol. 2014;88: 4366–4374. doi:10.1128/JVI.03134-13

109. Meade N, Furey C, Li H, Verma R, Chai Q, Rollins MG, et al. Poxviruses Evade Cytosolic Sensing through Disruption of an mTORC1-mTORC2 Regulatory Circuit. Cell. 2018;174: 1143–1157.e17. doi:10.1016/j.cell.2018.06.053

110. Meade N, King M, Munger J, Walsh D. mTOR Dysregulation by Vaccinia Virus F17 Controls Multiple Processes with Varying Roles in Infection. J Virol. 2019;93. doi:10.1128/JVI.00784-19

111. Meade N, Toreev HK, Chakrabarty RP, Hesser CR, Park C, Chandel NS, et al. The poxvirus F17 protein counteracts mitochondrially orchestrated antiviral responses. Nat Commun. 2023;14: 7889. doi:10.1038/s41467-023-43635-y

112. Luker KE, Hutchens M, Schultz T, Pekosz A, Luker GD. Bioluminescence imaging of vaccinia virus: effects of interferon on viral replication and spread. Virology. 2005;341: 284–300. doi:10.1016/j.virol.2005.06.049

113. Cureton DK, Massol RH, Saffarian S, Kirchhausen TL, Whelan SPJ. Vesicular stomatitis virus enters cells through vesicles incompletely coated with clathrin that depend upon actin for internalization. PLoS Pathog. 2009;5: e1000394. doi:10.1371/journal.ppat.1000394

114. Love MI, Huber W, Anders S. Moderated estimation of fold change and dispersion for RNA-seq data with DESeq2. Genome Biol. 2014;15: 550. doi:10.1186/s13059-014-0550-8

115. Lin Y-CJ, Li J, Irwin CR, Jenkins H, DeLange L, Evans DH. Vaccinia virus DNA ligase recruits cellular topoisomerase II to sites of viral replication and assembly. J Virol. 2008;82: 5922–5932. doi:10.1128/JVI.02723-07

