## Supplementary figures and images for "Metabolic reprogramming tips vaccinia virus infection outcomes by stabilizing interferon-γ induced IRF1"

### S1 Fig

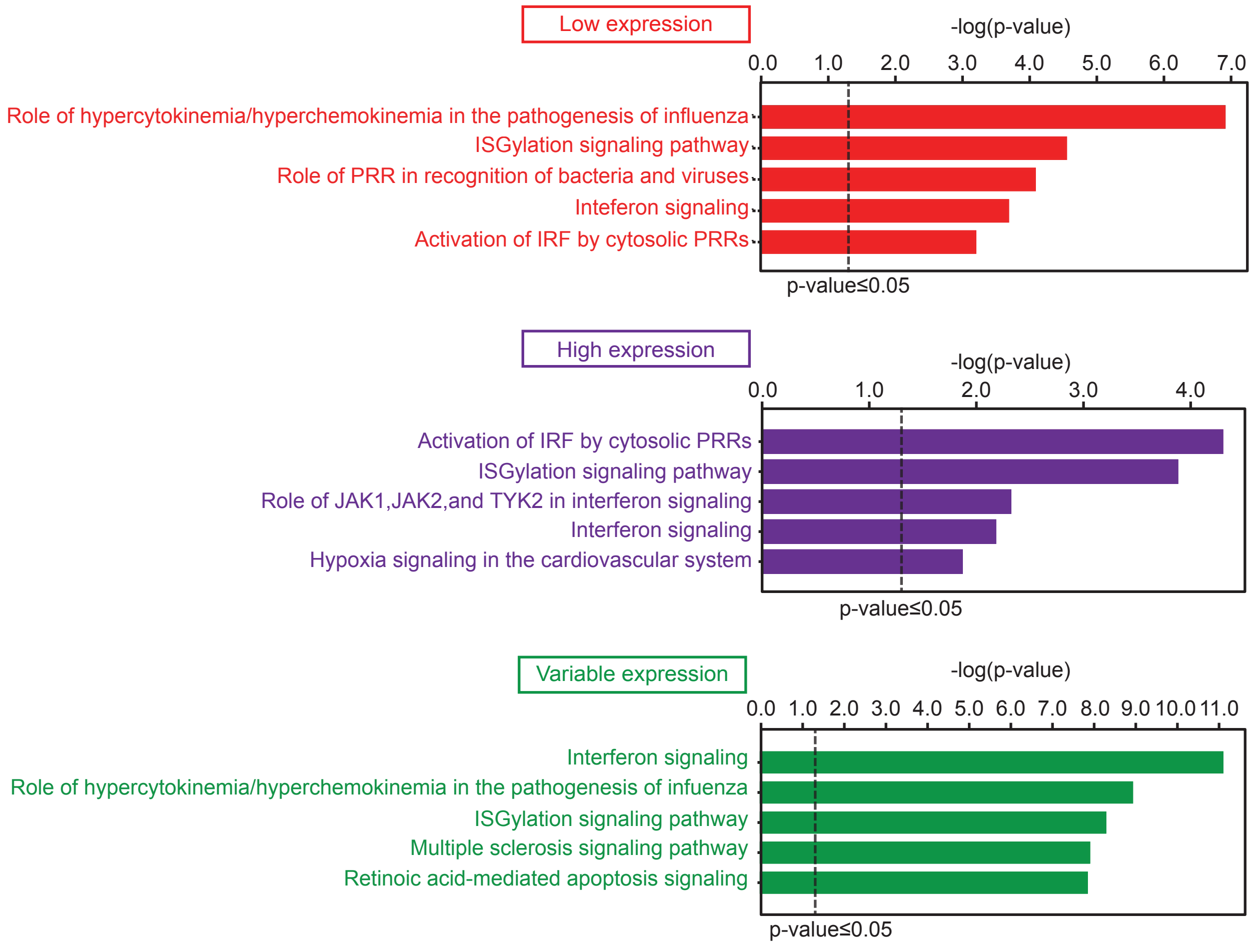

### S2 Fig

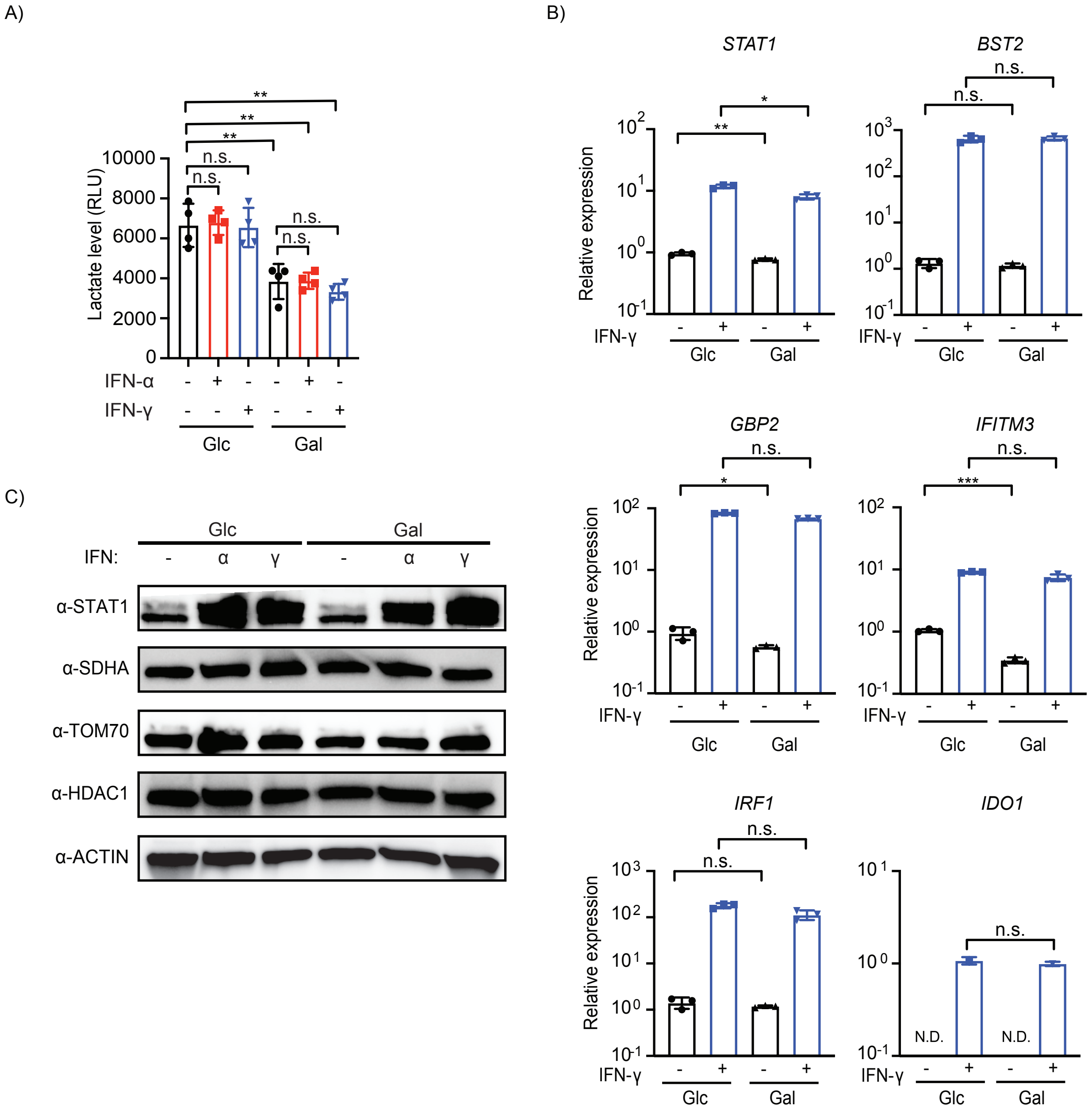

### S3 Fig

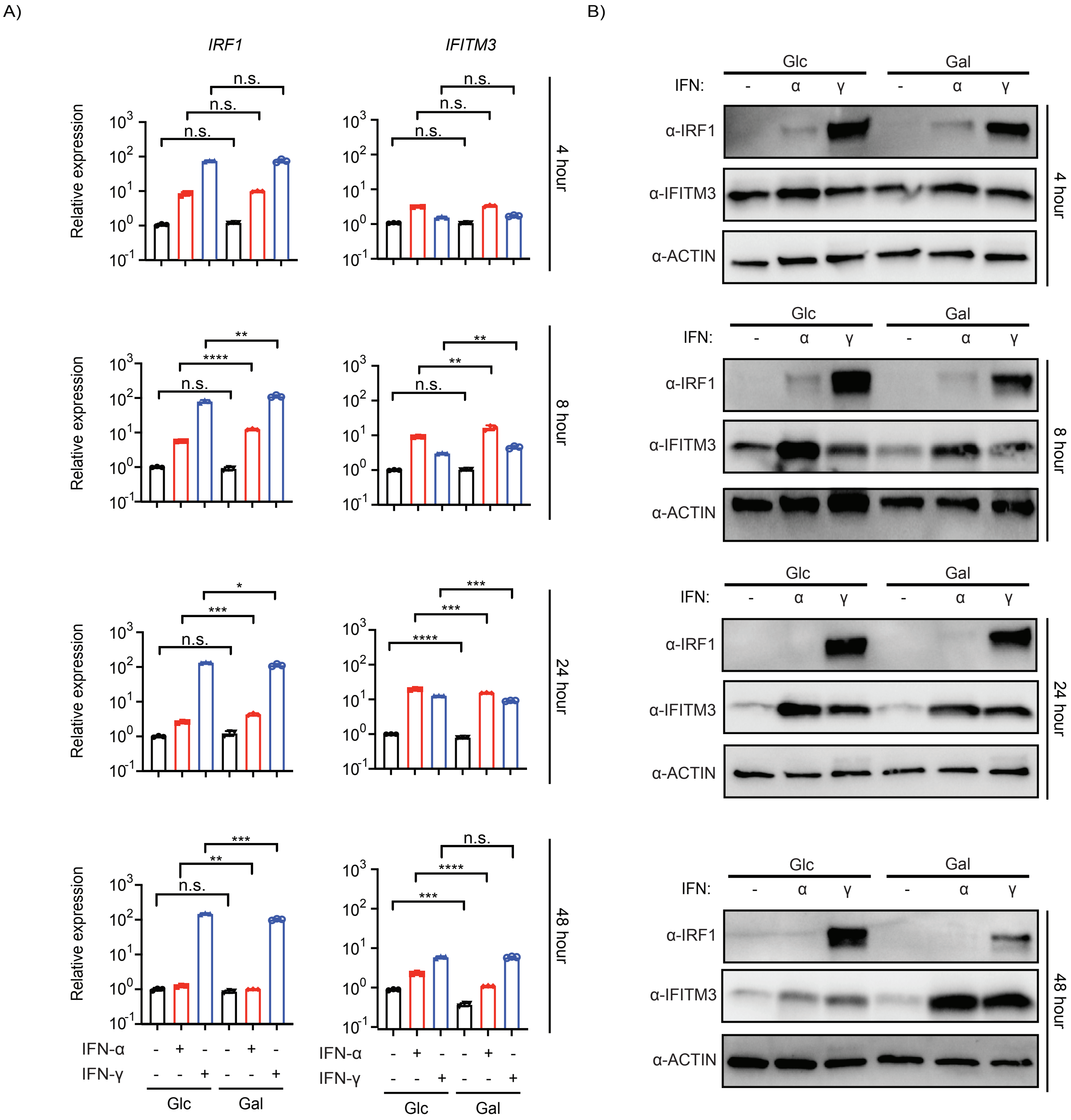

### S4 Fig

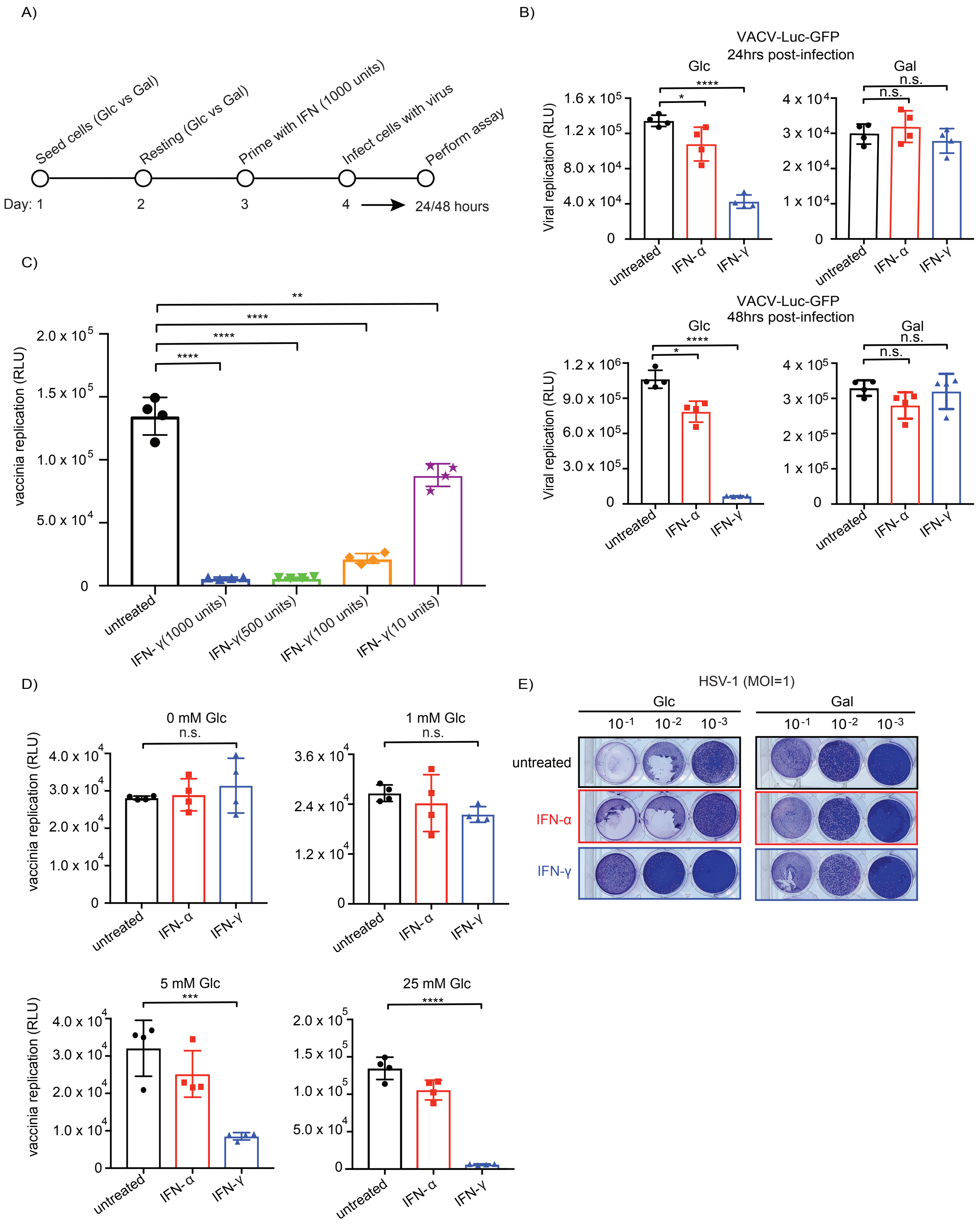

### S5 Fig

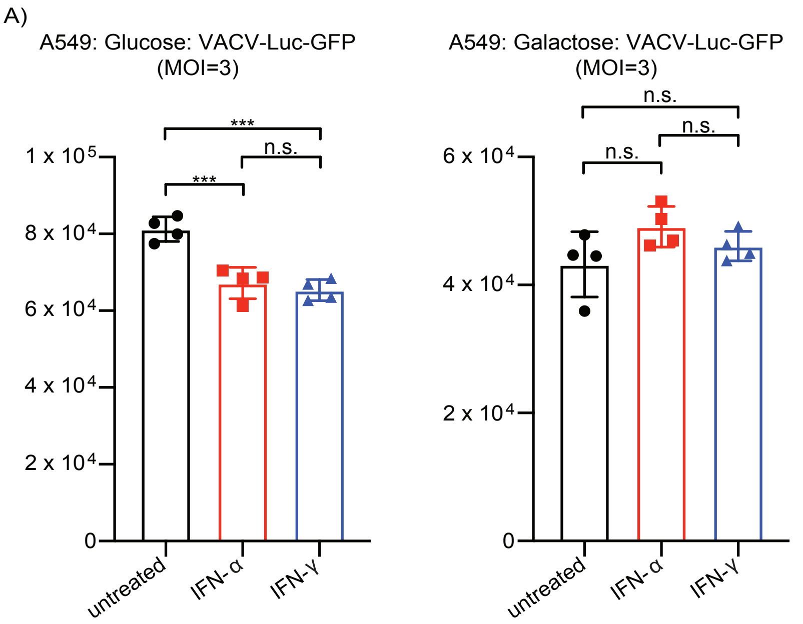

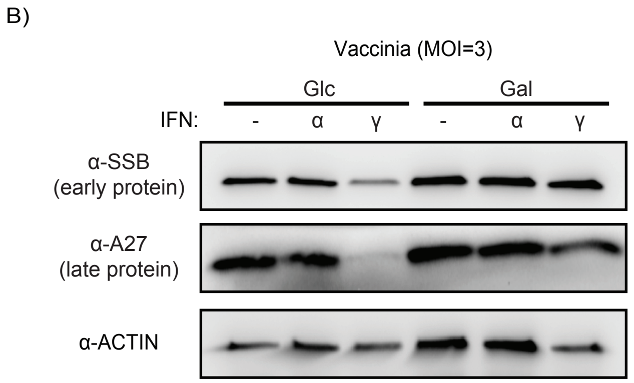

### S6 Fig

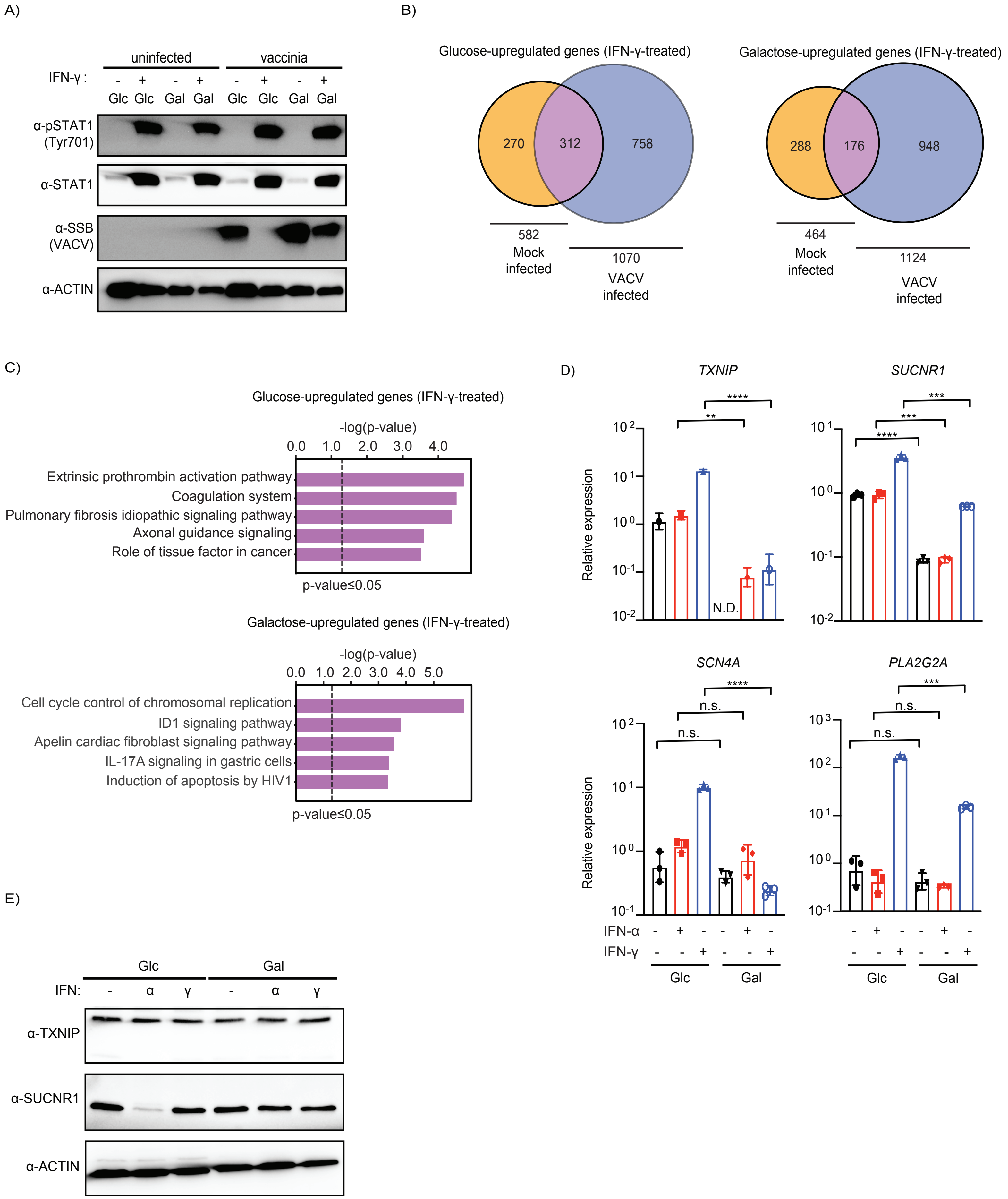

### S7 Fig

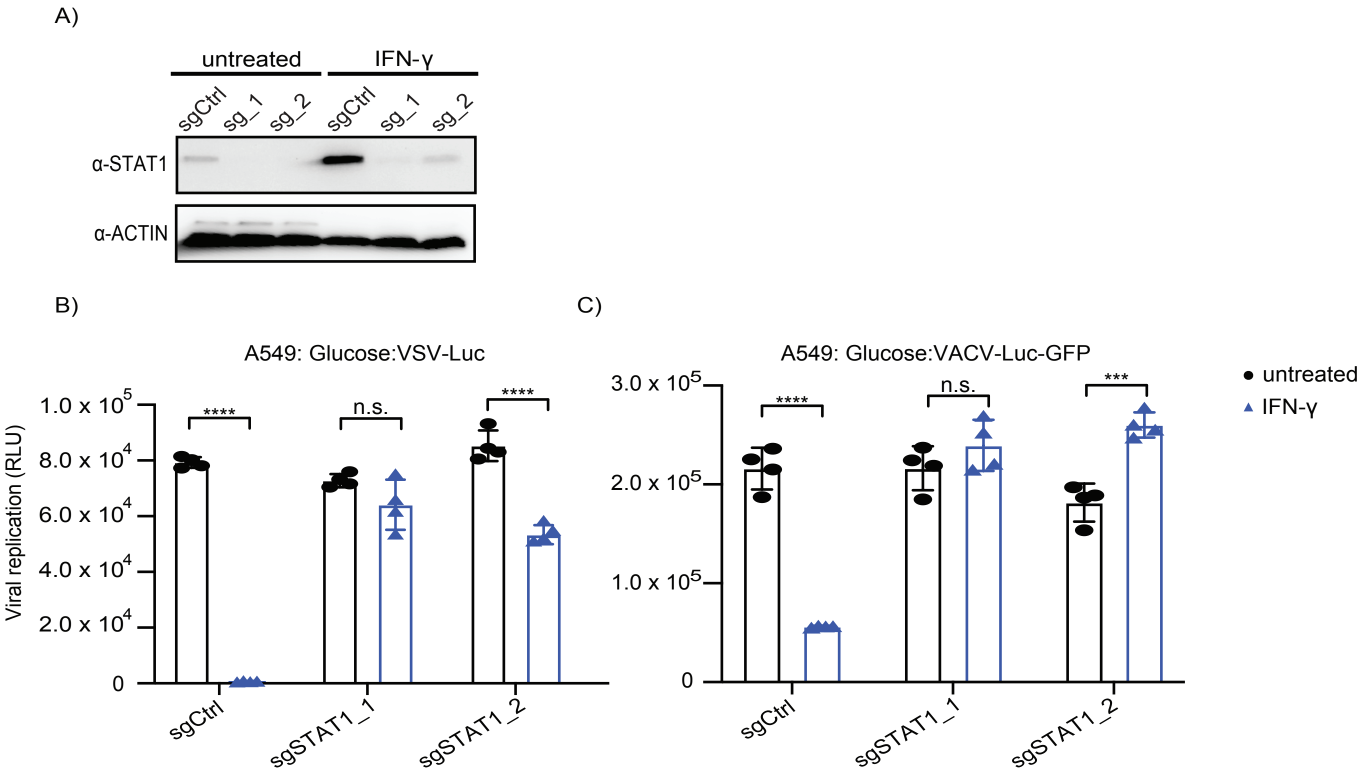

### S8 Fig

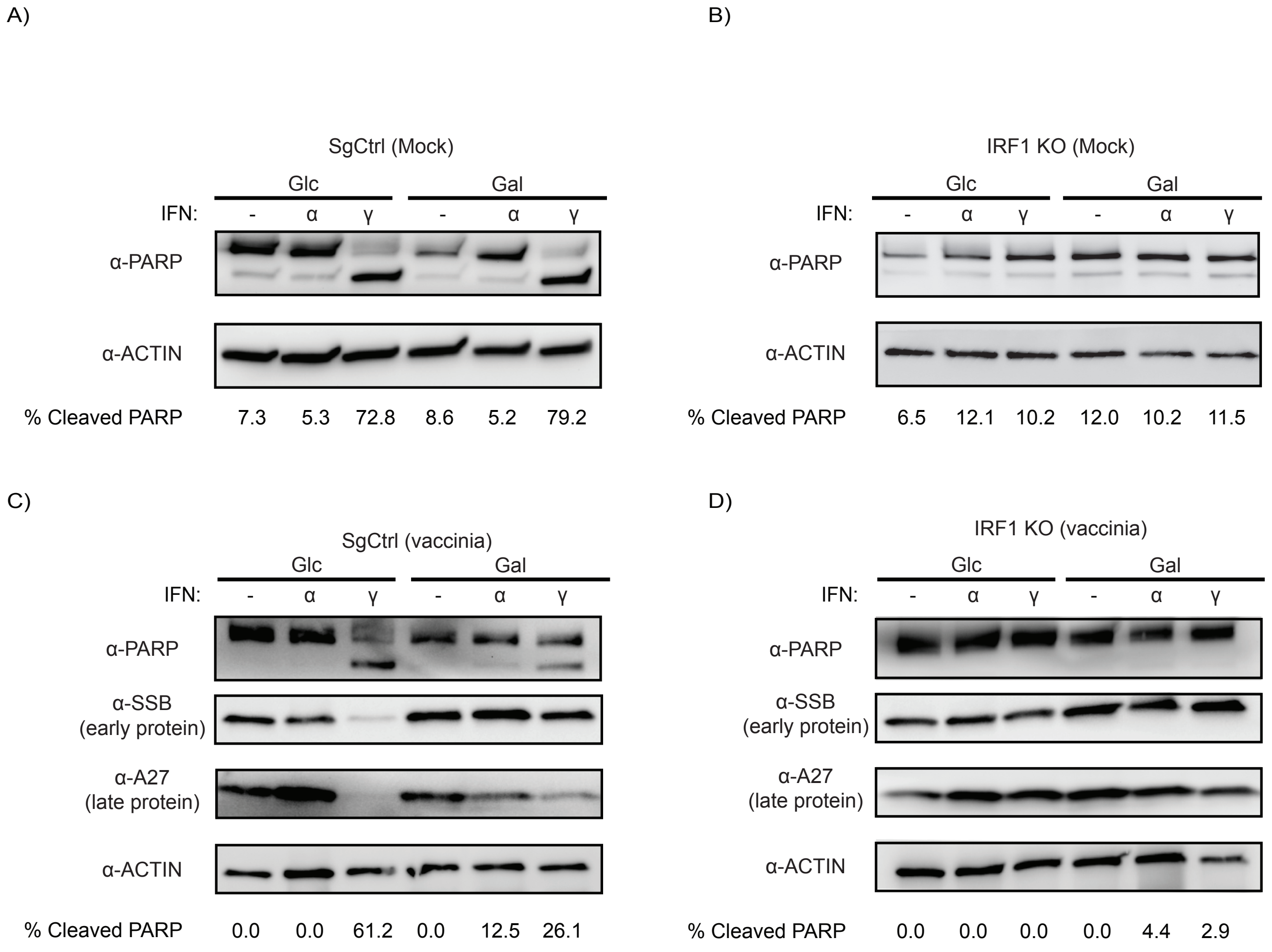

### S9 Fig

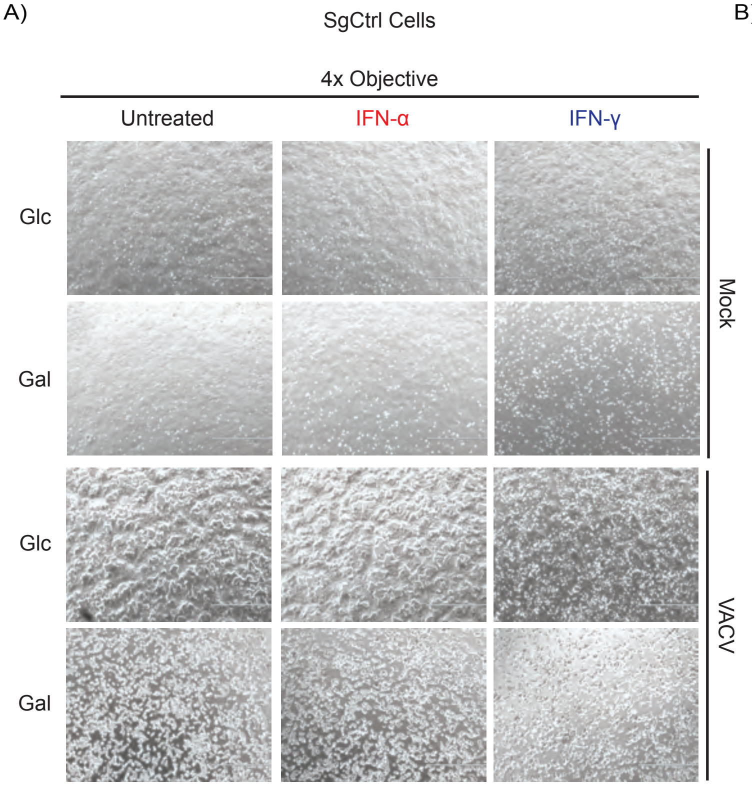

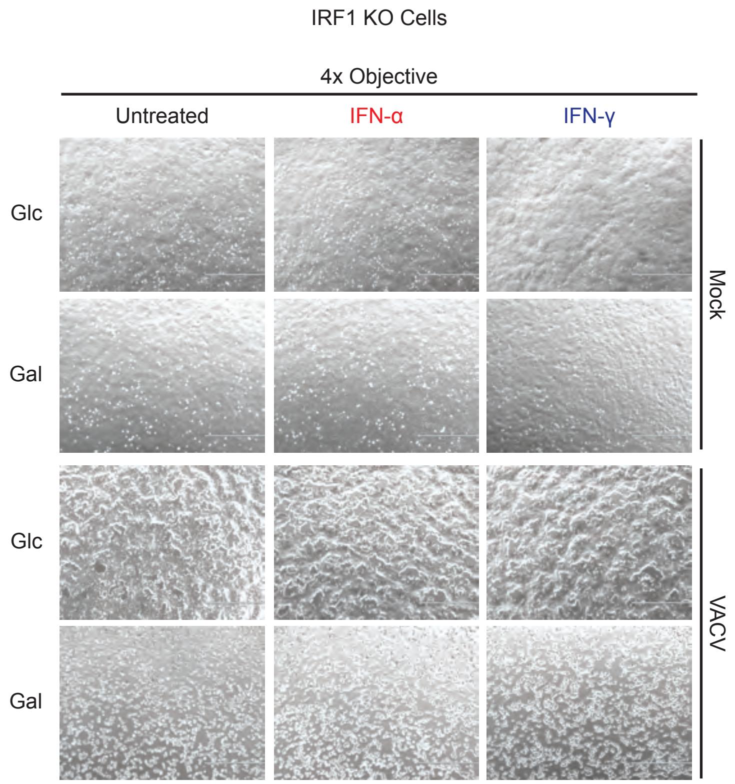

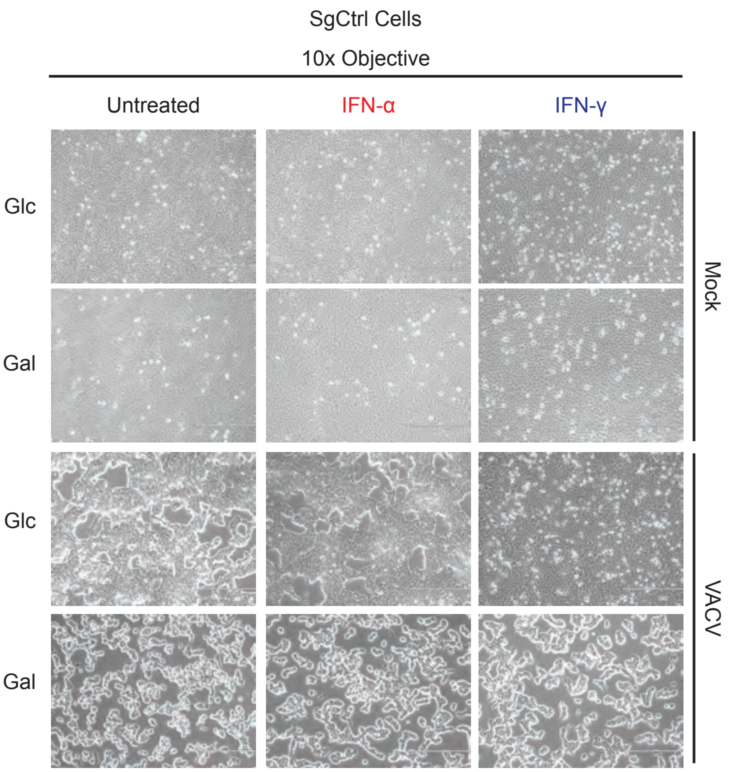

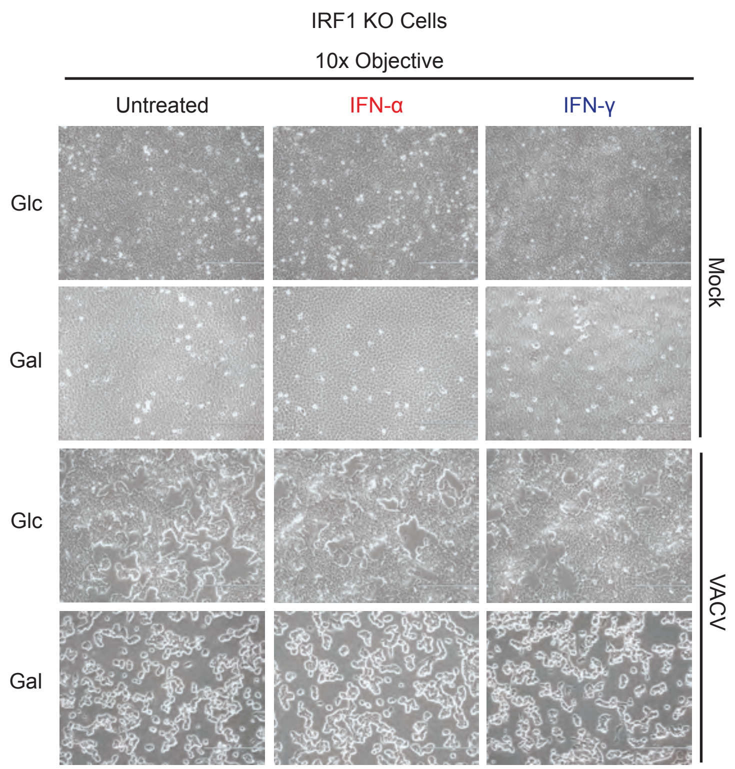

### S10 Fig

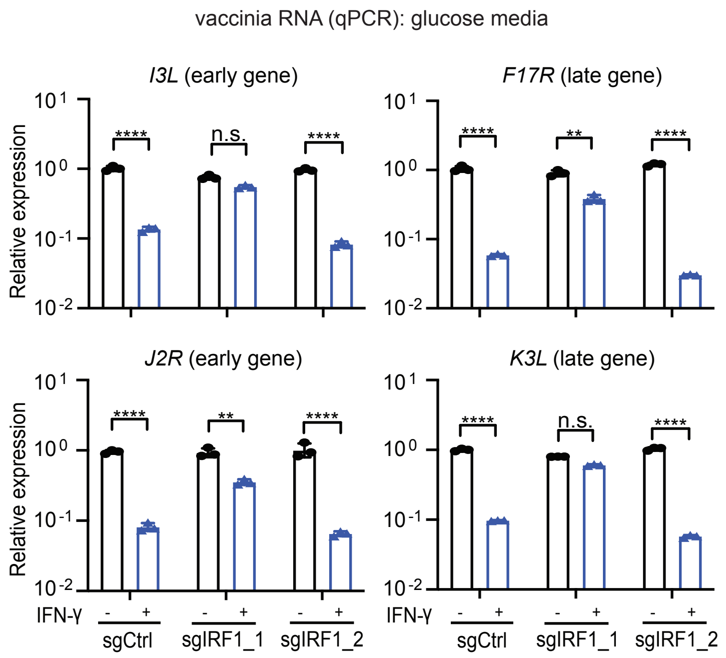

### S11 Fig

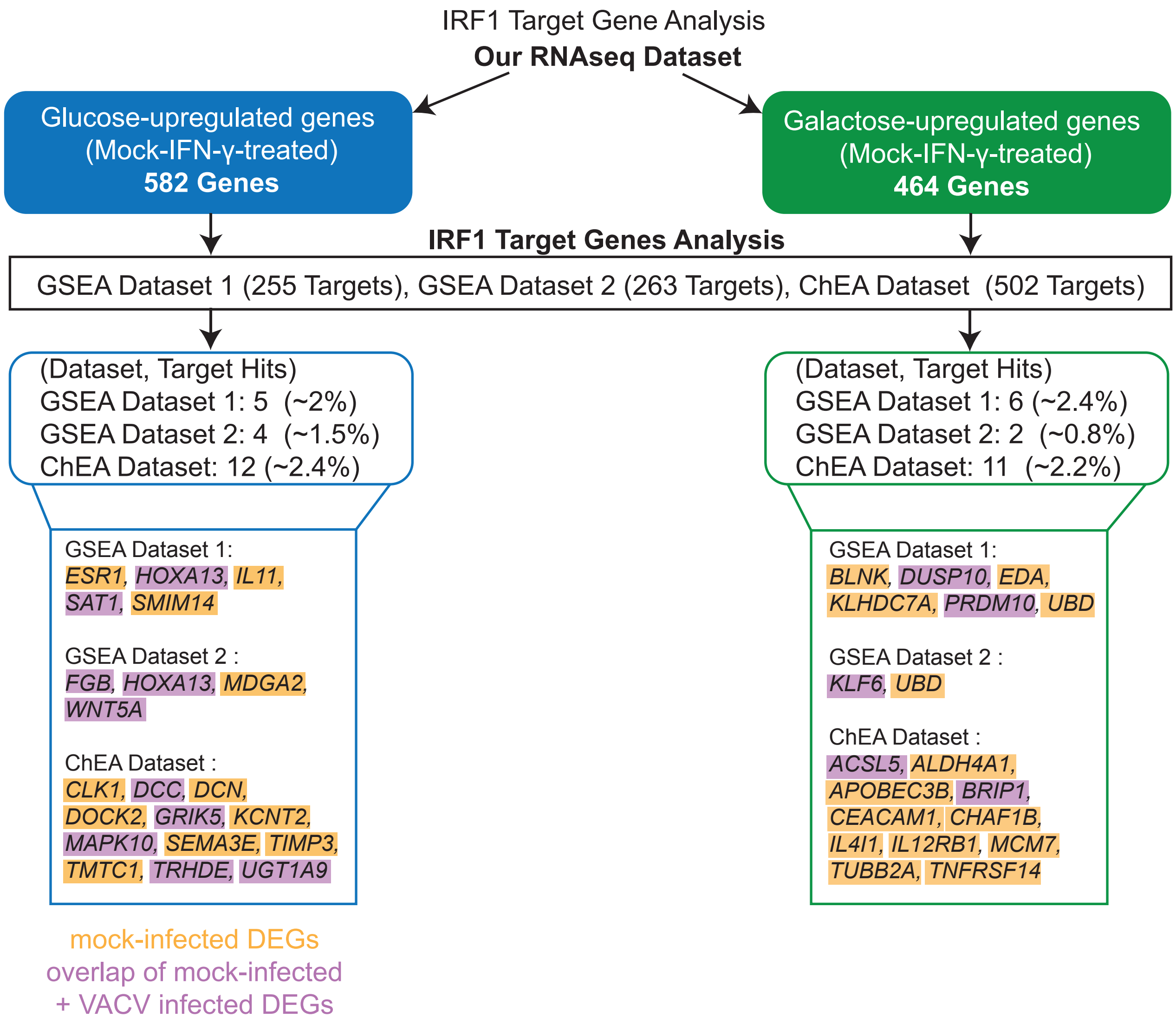
